# Single-cell information analysis reveals small intra- and large intercellular variations increase cellular information capacity

**DOI:** 10.1101/653832

**Authors:** Takumi Wada, Mitsutaka Wataya, Masashi Fujii, Ken-ichi Hironaka, Miki Eto, Shinsuke Uda, Daisuke Hoshino, Katsuyuki Kunida, Haruki Inoue, Hiroyuki Kubota, Hiroki Hamaguchi, Yasuro Furuichi, Yasuko Manabe, Nobuharu L. Fujii, Shinya Kuroda

## Abstract

Cells transmit information about extracellular stimulation through signaling pathways to control cellular function. A signaling pathway can be regarded as a communication channel. In the analysis of channels of cell populations, intercellular variation is considered noise. However, intercellular variation enables individual cells to encode different information. Therefore, at the single-cell level, each cell can be regarded as an independent channel. Thus, we propose that responses of cells of the same type in tissues, such as the fibers in a skeletal muscle, should be regarded as a multiple-cell channel composed of single-cell channels, in which intercellular variation contains information. Here, we applied electrical pulses to individual myotubes from cultured C2C12 cells or dissociated skeletal muscle fibers and measured Ca^2+^ responses or contraction, respectively, to estimate information capacity in a biological system. For each muscle cell system, we found that a single-cell channel transmitted more information than did a cell-population channel, indicating that the cellular response is consistent with each cell (low intracellular variation) but different among individual cells (high intercellular variation). As cell number and thus the number of single-cell channels increased, a multiple-cell channel transmitted more information by incorporating the differences among individual cells. Thus, a tissue with small intracellular and large intercellular variations has the capacity to distinguish differences in stimulation intensity to precisely control physiological function.

**One Sentence Summary:** Small intracellular and large intercellular variations increase information transmission for precise control of tissue function.

---

Signaling pathways transmit information about extracellular stimulation to control various cellular functions (*1*). These pathways must reliably convert stimulation intensity into signaling activity in the presence of noisy conditions, arising from noise within a cell (intracellular, intrinsic noise) and noise among cells (intercellular, extrinsic noise). An example of intracellular variation is the stochastic fluctuation of a biochemical reaction; examples of intercellular variation, are the differences in gene expression and protein abundance among cells (*2, 3*). The amount of information reliably transmitted through a signaling pathway can be quantified by mutual information (*4–21*) (Fig. 1, A and B). The mutual information corresponds to the logarithm of the average number of controllable states of cellular response that are defined by varied intensity of stimulation. The mutual information is determined by the balance between intensity and variation of a cellular response. The smaller the variation, the more information can be transmitted through a pathway with the same dynamic range. Even a response to high-intensity stimulation cannot be reliably transmitted if the variation is large (Fig. 1A, top). By contrast, even a response to a low-intensity stimulation can be reliably transmitted if the variation is small (Fig. 1A, bottom). Previous studies applying information theory to signaling pathways have examined the cell-population level (*4–20*) (table S1), in which a cell population, but not single cells, are regarded as a single communication channel [defined hereafter as a cell-population channel (Fig. 1C, left)]. For a cell-population channel, the mutual information relating stimulation intensity and cellular response contains only 1 bit information (Fig. 1C, left, table S1) (*4–20*), indicating that the cellular response can distinguish only the two conditions of stimulation: the presence or absence of stimulation. Physiologically, cellular responses are often gradually controlled by the intensity of stimulation, meaning that a cellular response can encode more than 1 bit information.

**Fig. 1.**
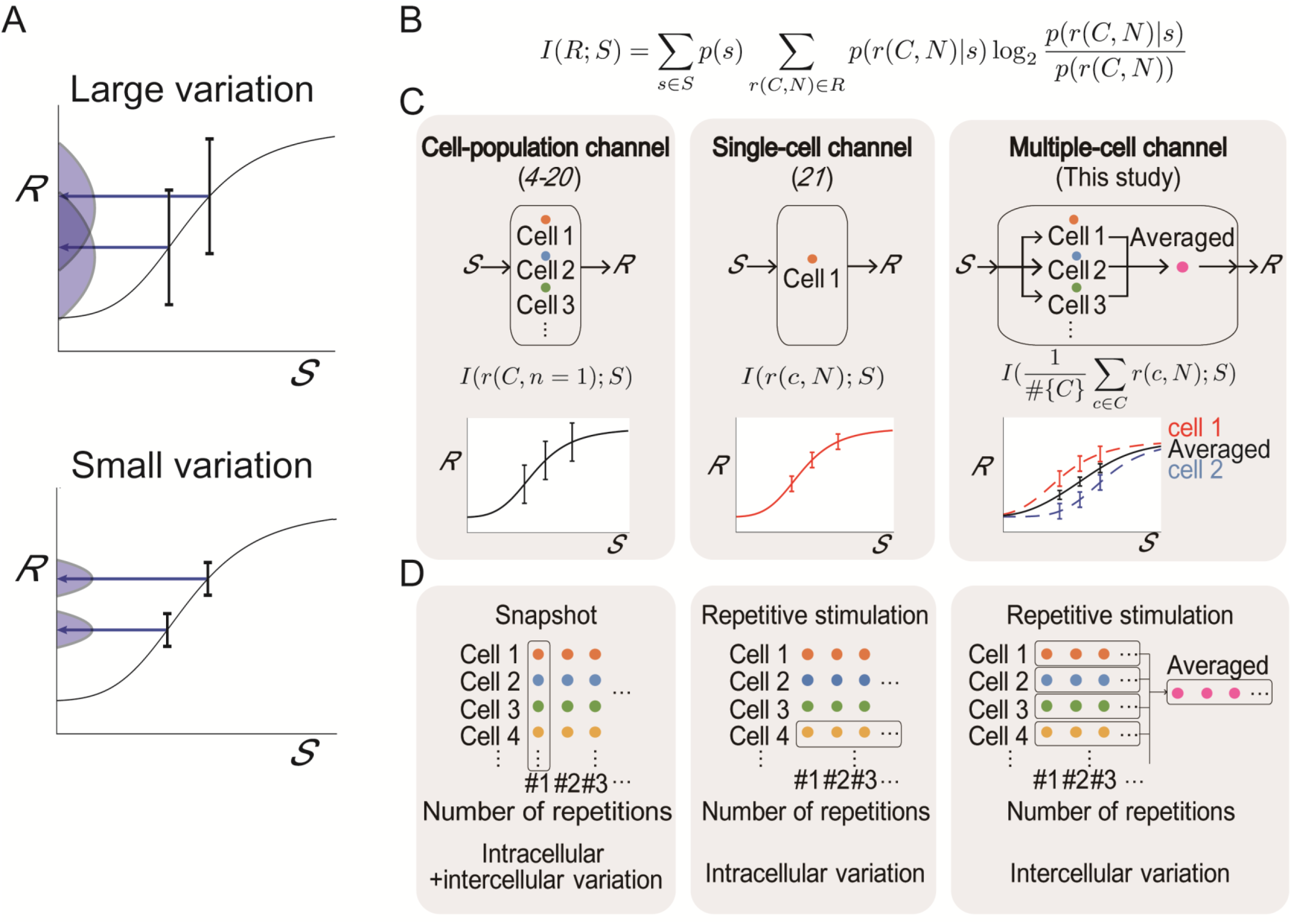
Information transmission in a cell-population, a single-cell, and a multiple-cell channel. (**A**) The relationship between stimulation (*S*) and response (*R*) and dependence of the ability to discriminate between high and low intensity stimulation on response variation. (**B**) Mutual information between stimulation and response. *S, R, C,* and *N* represent the set of stimulations, responses, cells, and the number of repetitions of stimulation, respectively. *p*(*S*) and *p*(*r*(*C, N*)) are probability distributions of stimulation intensity and response intensity, respectively, and *p*(*r*(*C, N*)|*S*) is a conditional probability distribution of the response for a given stimulation. (**C**) Mutual information of a cell-population, a single-cell, and a multiple-cell channel. For a cell-population channel, response probability distribution is calculated from a single stimulation for each cell and includes both intracellular and intercellular variations as noise. For a single-cell channel, response probability distribution is calculated from repetitive stimulation for each cell and includes intracellular variation as information. A multiple-cell channel, composed of a combination of single-cell channels, includes both intercellular and intracellular variation as information. Mutual information in the different channels differs in the definition of response *R*, *r*(*C, n* = 1), *r*(*c, N*), 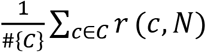 for a cell-population, single-cell channel and multiple-cell channel respectively, where *n* ∈ *N* is the number of repetitions of stimulation, and *c* ∈ *C* is the cell.

One of the limitations in analyzing a biological system as a cell-population channel is that intercellular variation is regarded as noise or uncertainty of signal intensity. However, each cell can be regarded as an independent single-cell channel, because intercellular variation enables individual single-cell channels to encode different information (Fig. 1C, middle). In some tissues and organs, such as skeletal muscle, the response represents the sum of responses of individual cells. Therefore, we propose that organs and tissues are better represented by a communication channel composed of a sum of single-cell channels, which we define as a multiple-cell channel (Fig. 1C, right). We use the average of the outputs of single-cell channels, rather than a sum of the single-channel outputs, to determine the output of a multiple-cell channel so that we can directly compare single-cell channels with multiple-cell channels (Fig. 1, C and D). In a multiple-cell channel, each single-cell channel can encode different information, and intercellular variation represents information. Therefore, the analysis of single-cell channels and the incorporation of these into a multiple-cell channel will accurately quantify how much information is physiologically transmitted in a tissue or organ (Fig. 1C).

Measuring the mutual information in a single-cell channel requires acquiring data of cellular response (*R*) to repetitive stimulation (*S*) with various intensities for individual cells (Fig. 1C, middle). These data enable to calculate the probability distribution of *R* for a given *S*, *p*(*R*|*S*) and the mutual information in a single-cell channel, *I*(*R*;*S*) (Fig. 1B, C, middle). For some experimental paradigms, repetitive stimulation with various intensities of stimulation takes a long time, leading to change in the abundance of molecules and a change in an internal state of a cell. Thus, the cellular response varies with time, as well as stimulation intensity, complicating the calculation of the mutual information between stimulation and cellular response. Analysis of a single-cell channel from the Ca^2+^ response in HEK293 cells stimulated with acetylcholine repetitively at various concentrations enabled the calculation of the channel capacity, which is the maximal amount of information that the channel can transfer, at 2 bits (*21*). This 2-bit channel capacity is higher than the 1-bit capacity calculated for cell-population channels in other studies (*4–20*). With organs, such as skeletal muscle, functioning not as a single cell, but as the sum of single cells, a multiple-cell channel should correspond to the physiological communication channel. Because intercellular variation enables individual single-cell channels to encode different information, a multiple-cell channel is likely to transmit more information by incorporating intercellular variation as information. However, how much information a multiple-cell channel can transfer has not been explored.

In this study, we solved a technical problem of avoiding time variance in response to repetitive stimulation by measuring Ca^2+^ response induced by repetitive electrical pulse stimulation (EPS) in differentiated C2C12 myotubes (*22*) (figs. S1 and S2). We established C2C12 cell lines stably expressing GCaMP6f, a fluorescent Ca^2+^ probe (*23*), and differentiated the cells into myotubes. In C2C12 myotubes, the Ca^2+^ response to a single EPS lasted ~1 second, and Ca^2+^ amplitude was consistent for 200 repetitive EPS (fig. S2). Thus, the Ca^2+^ response is a time-invariant response, enabling calculation of the mutual information. To calculate the mutual information between EPS and Ca^2+^ response in a single-cell channel of a C2C12 myotube, we repetitively stimulated each myotube 20 times at 10 different voltages and measured the amplitude of the Ca^2+^ response. We established that the 10 voltages were sufficient to avoid underestimation of mutual information (fig. S3) and that, although an overestimation bias was present, it was small enough to have little effect on the calculation of the mutual information from data acquired with 20 repetitive stimulations of 551 individual myotubes (fig. S4).

From the analysis of 551 C2C12 myotubes, we observed that Ca^2+^ amplitude generally increased as the voltage increased (Fig. 2A); however, even for the same voltage, the Ca^2+^ amplitude varied from myotube to myotube (Fig. 2B). The variation of Ca^2+^ amplitude in each myotube was much smaller than that in the population (Total) (Fig. 2, B and C). The larger variation of the population derived from large intercellular variation (Fig. 2, D and E, Eqs. 2 and 3 in Materials and Methods). On average, intercellular variation accounted for 83% of the total variation, indicating that intercellular variation is much larger than intracellular variation.

**Fig. 2.**
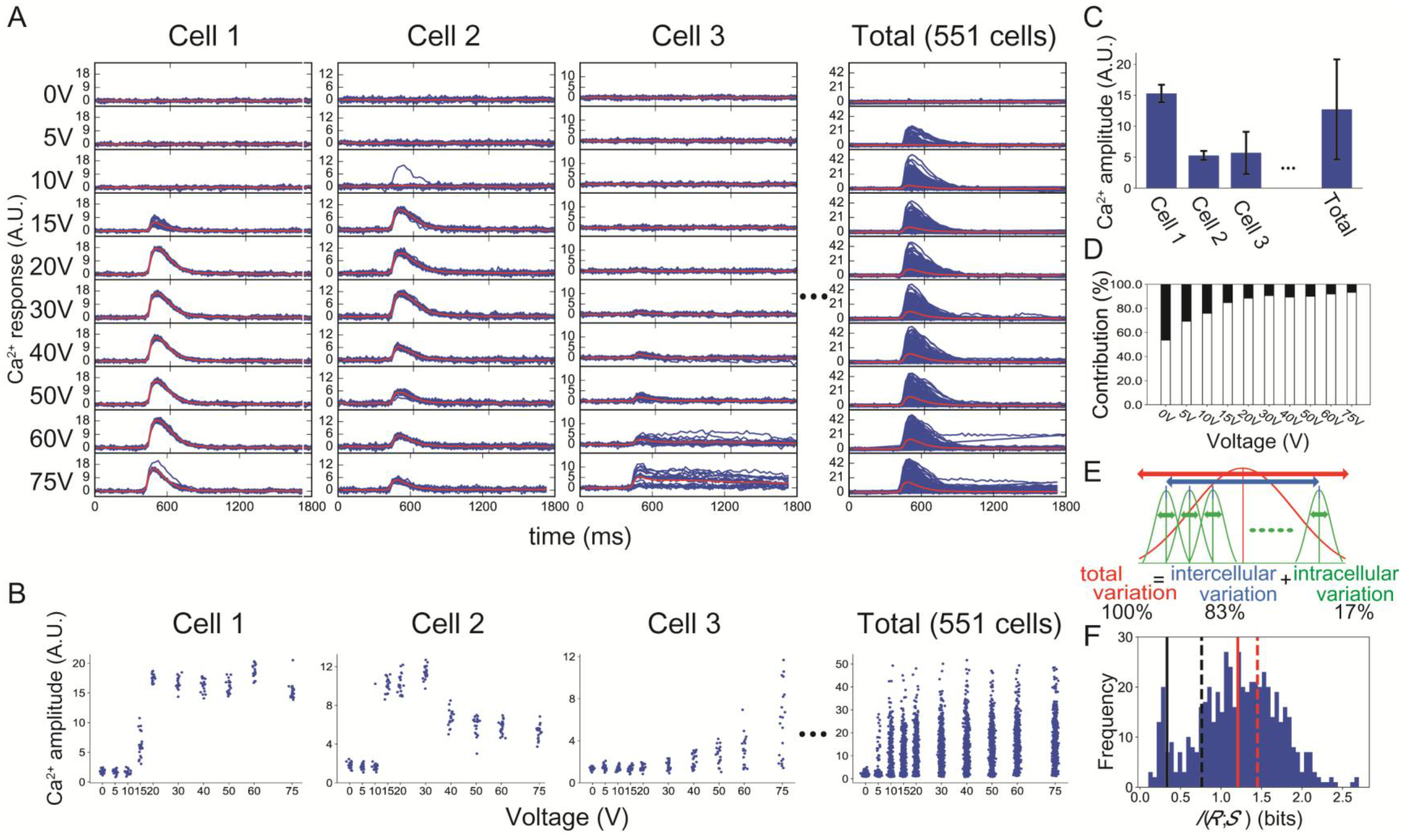
Information transmission of a single-cell channel for electrical pulse stimulation into increases in Ca^2+^ amplitude in C2C12 myotubes. (**A**) Ca^2+^ response in individual differentiated C2C12 myotubes by repetitive stimulation (20 times) with 10 different intensities of electrical pulse stimulation (EPS). “Cell 1” to “Cell 3” are representative single-cell responses. “Total” indicates the responses in 551 myotubes. Each blue line indicates the Ca^2+^ response induced by single stimulation for each cell. Red lines indicate the averaged response time course of the total cells. (**B**) Ca^2+^ amplitude versus voltage from the 3 cells shown in (A). A dot indicates Ca^2+^ amplitude to each single stimulation. Ca^2+^ amplitude is defined as the maximum response subtracted by basal Ca^2+^ before stimulation (fig. S2). (**C**) Average Ca^2+^ amplitudes induced by 75V EPS in individual C2C12 myotubes and in the total population. The error bars indicate standard deviations. (**D**) The percentages of intercellular (white) and intracellular (black) variation in the total variation for the indicated voltage of EPS (Eqs. 2 and 3 in Materials and Methods). (**E**) The total variation (red) divided into intercellular (blue) and intracellular (green) variations. (**F**) Histogram of the mutual information between intensity of EPS and Ca^2+^ amplitude in single-cell channels. Red dashed line, the average of channel capacity of a single-cell channel (1.45 bits); red solid line, the average mutual information of a single-cell channel (1.21 bits); black dashed line, channel capacity of the cell–population channel (0.76 bits); black solid line, the mutual information of the cell–population channel (0.33 bits). We used the optimal input probability distribution for the averaged response of the total cells to calculate the mutual information, and the optimal input probability distribution for each channel to calculate channel capacity.

To quantify how much information about voltage can be encoded in a Ca^2+^ response (amplitude) in a single-cell channel and cell-population, we calculated the probability distribution of Ca^2+^ amplitudes at each voltage for each myotube, calculated the mutual information of each single-cell channel relating voltage with Ca^2+^ amplitude, then plotted the frequency of the mutual information values occurred for the 551 myotubes (Fig. 2F). Calculation of the mutual information requires an input probability distribution, which cannot be determined for a biological system. Thus, we calculated the mutual information using optimal input probability distribution for the averaged response of the total myotubes (see Materials and Methods). We assumed that muscle function (fiber contraction) represents the sum of the function of the myotubes comprising it; thus, muscle function is controlled by the same optimal input probability distribution as that controlling the individual myotubes. To define this input, we calculated the averaged response of 551 myotubes and estimated the optimal input probability distribution of the averaged response for the total (fig. S5, A and B). We used this optimal input probability distribution to the averaged response to calculate mutual information in a single-cell channel for each myotube. The average mutual information in a single-cell channel was 1.21 ± 0.50 bits (mean ± S.D.) (Fig. 2F, red solid line). Note that *n*-bit indicates 2^*n*^ states. The result indicates that among the 10 voltage conditions (3.32 bits), on average a single-cell can distinguish 2.32 conditions. Using the optimal input probability distribution for the averaged response of the total myotubes, we calculated the mutual information in the cell-population channel as 0.33 bits (Fig. 2F, black solid line), indicating that on average the cell-population channel cannot distinguish between 2 conditions. This means that as a population, the myotubes cannot distinguish even the presence or absence of stimulation. Thus, the single-cell channel transmitted more information than the cell-population channel, because the cell-population channel includes the high intercellular variation.

Using the optimal input probability distribution for each single-cell channel, we calculated the channel capacity, which is the maximum amount of information that can be transmitted, of the single-cell channel as 1.5 bits (Fig 2F, red dashed line). This value was similar to and within the standard deviation of the average channel capacity. Furthermore, there was a strong correlation (0.947) between mutual information with the optimal input probability distribution for the averaged response and channel capacity of single-cell channels (fig. S6). Therefore, hereafter unless otherwise specified, we defined the “mutual information” as the mutual information calculated with the optimal input probability distribution for the average response of the total cells, and we defined “channel capacity” as the mutual information calculated with the optimal input probability distribution for each channel. The channel capacity of the myotube cell-population was similar to those of previous studies of cell-population channels (~1 bit) (table S1) (*4–20*). We obtained similar results for the mutual information and channel capacities for single-cell channels and cell-population channels in two independent clones of C2C12 myotube lines stably expressing GCaMP6f (fig. S7A and B). Additionally, these two clones also exhibited greater intercellular variation than intracellular variation (fig. S7C and D).

In the cell-population channel, intercellular variation is regarded as uncertainty of signal intensity and represents noise. Therefore, the mutual information in the cell-population channel is lower than that in the single-cell channel. In contrast, intercellular variation enables individual single-cell channels to encode different information, such as different signal intensities, suggesting that a multiple-cell channel composed of a sum of single-cell channels can encode more information than a single-cell channel. To examine this possibility, we calculated the mutual information of a 2-cell channel for every pair of myotubes (Fig. 3). The 2-cell channel is a sum of two different single-cell channels (Eq. 10 in Materials and Methods). We averaged the responses of two paired cells to calculate the probability distribution of the averaged response, which we used as the response of a 2-cell channel to calculate the mutual information of a 2-cell channel. The average mutual information of the 2-cell channel was 1.55 ± 0.43 bits (mean ± S.D.) (Fig. 3A). This result indicates that on average 2.94 conditions can be distinguished by a 2-cell channel and that a 2-cell channel can transmit more information and distinguish more conditions than a single-cell channel, which only distinguished 2.32 conditions on average. Moreover, increasing the number of cells in a multiple-cell channel increased the mutual information (Fig. 3B, blue). In the myotube experiments with 10 stimulation intensities, the mutual information plateaued with the inclusion of ~2^7^ myotubes at 3.13 bits, because the mutual information approached the amount of information of input (3.32 bit), suggesting that the mutual information for a multiple cell channel with greater than 2^7^ cells is underestimated.

**Fig. 3.**
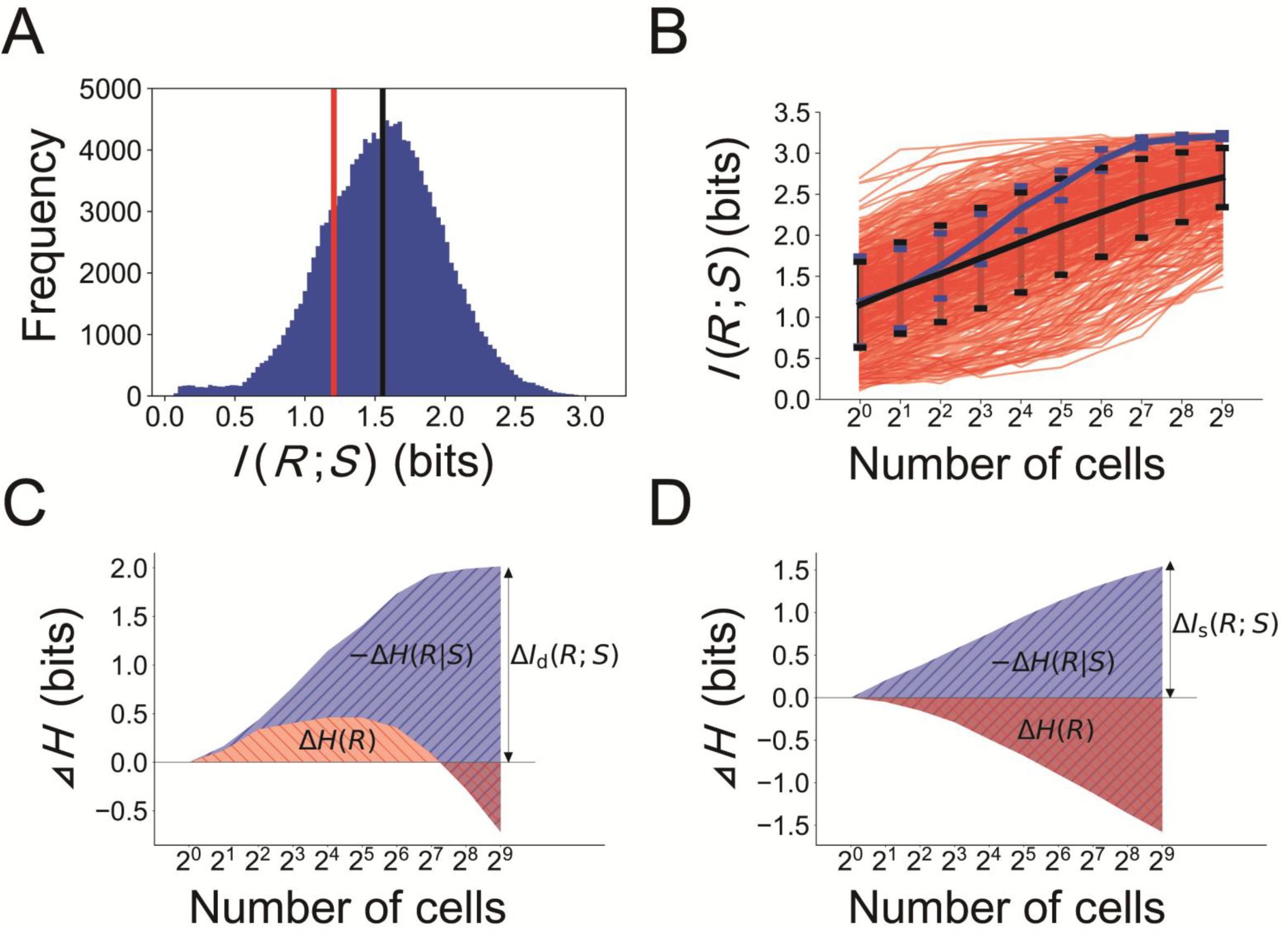
Information transmission of a multiple-cell channel composed of single-cell channels. (**A**) Histogram of the mutual information of a multiple-cell channel comprised of 2-cell channels (Eq. 10 in Materials and Methods). Black line, the average mutual information of 2-cell channels (1.55 bits); red line, the average mutual information of single-cell channels (1.21 bits). (**B**) Mutual information of a multiple-cell channel according to the number of cells included as single-cell channels. Blue line, the average mutual information of a multiple-cell channel composed of single-cell channels from the 551 different myotubes, defined as *I*_d_(*R*; *S*). Red lines, mutual information of a multiple-cell channel composed of the same myotube by resampling responses repetitively from the same myotube, defined as *I*_s_(*R*; *S*). Black line, the average of *I*_s_(*R*; *S*). Bars indicate standard deviation for both black and blue lines. (**C**) The contribution of the difference in the average *H*(*R*), Δ*H*(*R*), and the difference in the average −*H*(*R*|*S*), −Δ*H*(*R*|*S*), to differences in the average *I*_d_(*R*; *S*), Δ*I*_d_(*R*; *S*) in a multiple-cell channel composed of different single-cell channels. The differences were defined by those between *H*(*R*), −*H*(*R*|*S*), *I*_d_(*R*; *S*) of each number of cells and those whose size is 1 (Eqs. 15, 16, and 17 in Materials and Methods). (**D**) The contribution of Δ*H*(*R*) and −Δ*H*(*R*|*S*) to Δ*I*_s_(*R*; *S*) in a multiple-cell channel composed of the same single-cell channel. We used the optimal input probability distribution for the averaged response of the total cells to calculate the mutual information.

To examine the contributions of intracellular and intercellular variations to the increase in the mutual information, we virtually created a multiple-cell channel composed of the same myotube by resampling responses repetitively from the same cell, where intercellular variation is not involved. The mutual information of a multiple-cell channel composed of the same myotube was calculated for 551 myotubes (Fig. 3B, red). Hereafter, we define the mutual information of a multiple-cell channel composed of different myotubes as *I*_d_(*R*; *S*) and that of the same myotube as *I*_s_(*R*; *S*). For a 2-cell channel, about half of *I*_s_(*R*; *S*) were larger than the average of *I*_d_(*R*; *S*) (compare number of red lines with blue value at 2^1^ on x-axis in Fig. 3B). However, as the number of myotubes in a multiple-cell channel increased, the average of *I*_d_(*R*; *S*) (Fig. 3B, blue) gradually became exceeded most of *I*_s_(*R*; *S*) (Fig. 3B, red). Because *I*_d_(*R*; *S*) includes intercellular variation, this result suggested that intercellular variation is a key component for information encoding of a multiple-cell channel. Then, we investigated why the mutual information of a multiple-cell channel was larger than that of a single-cell channel. Mutual information can be expressed as

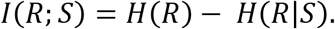

*H*(*R*) is the amount of information of the response; *H*(*R*) is illustrated as the average logarithm of the number of distinguishable states of responses. *H*(*R*|*S*) is the amount of information of the response for a given stimulation intensity; *H*(*R*|*S*) is illustrated as the average logarithm of the number of response for a given stimulation intensity. The mutual information becomes larger as *H*(*R*) becomes larger or *H*(*R*|*S*) becomes smaller. For a pair of identical cells, *H*(*R*) does not increase, indicating that any increase in *H*(*R*) depends on intercellular variation (fig. S8). For example, for a pair of cells with different responses to a set of stimulation intensities, *H*(*R*) can increase because of an increase in the distinguishable number of responses determined by averaging the responses of both cells (fig. S8A). At the same time, averaging the responses decreases *H*(*R*|*S*) because the variance of averaged response decreases as the number of cells increases. In contrast, for a pair of cells with identical responses to a set of stimulation intensities, averaging the responses decreases *H*(*R*|*S*) without any change in *H*(*R*) (fig. S8B), because there is no intercellular variation. In this case, intercellular variation does not contribute to the increase in the mutual information.

We examined the contribution of the change in Δ*H*(*R*) from the effect of intercellular variation and the change in −Δ*H*(*R*|*S*) to the increase in the mutual information of a multiple-cell channel (Fig. 3, C and D, Eqs. 13, 14, 16, 17 in Materials and Methods). Both Δ*H*(*R*) and −Δ*H*(*R*|*S*) contributed to Δ*I*_d_(*R*; *S*) (Fig. 3C). The rate of the increase in *I*_d_(*R*; *S*) against the increase in myotube numbers was larger than that of the increase in *I*_s_(*R*; *S*) from 2^2^ to 2^6^ myotubes, because of increase in Δ*H*(*R*). By contrast, only −Δ*H*(*R*|*S*) contributed to Δ*I*_s_(*R*; *S*) (Fig. 3D). The decrease Δ*H*(*R*) that occurred for multiple-cell channels composed of more than 2^5^ cells is an artifact caused by underestimation of Δ*H*(*R*) as a result of using discrete input probability distribution (fig. S9). If the input probability distribution is continuous, Δ*H*(*R*) increases monotonically as the number of cells increases (fig. S9). Thus, with a continuous input probability distribution, Δ*H*(*R*) would contribute to the increase in the mutual information for the entire range. Because the increase in Δ*H*(*R*) is due to the intercellular variation, these results indicate that intercellular variation increases the information capacity of a multiple-cell channel. Cheong *et al* (*12*) showed an increase in the information capacity of a multiple channel composed of cell-population channels. Thus, increase the number of channels incorporated into a multiple channel based on either cell-population channels or single-cell channels increased information capacity.

The Ca^2+^ response triggers skeletal muscle contraction, representing a final biological output. However, in C2C12 myotube, contraction is so weak that it is difficult to acquire the quantitative contraction data. Therefore, we isolated single fibers from flexor digitorum brevis (FDB) muscle of mice and used EPS-induced contraction of single fibers as a physiological output (*28*). We measured contraction in response to 10 repetitive stimulations at 32 different voltages and calculated the mutual information between EPS and contraction (Fig. 4, fig. S10) (Materials and Methods). Even for the same voltage, contraction varied from fiber to fiber (fig. S10). The variation of contraction in each fiber was much smaller than that across the population (fig. S10B). The larger variation of the fiber population derived from a large intercellular variation (Fig. 4A, Eqs. 2 and 3 in Materials and Methods). Similar to the C2C12 myotubes, on average, intercellular variation accounted for 86% of the total variation.

**Fig. 4.**
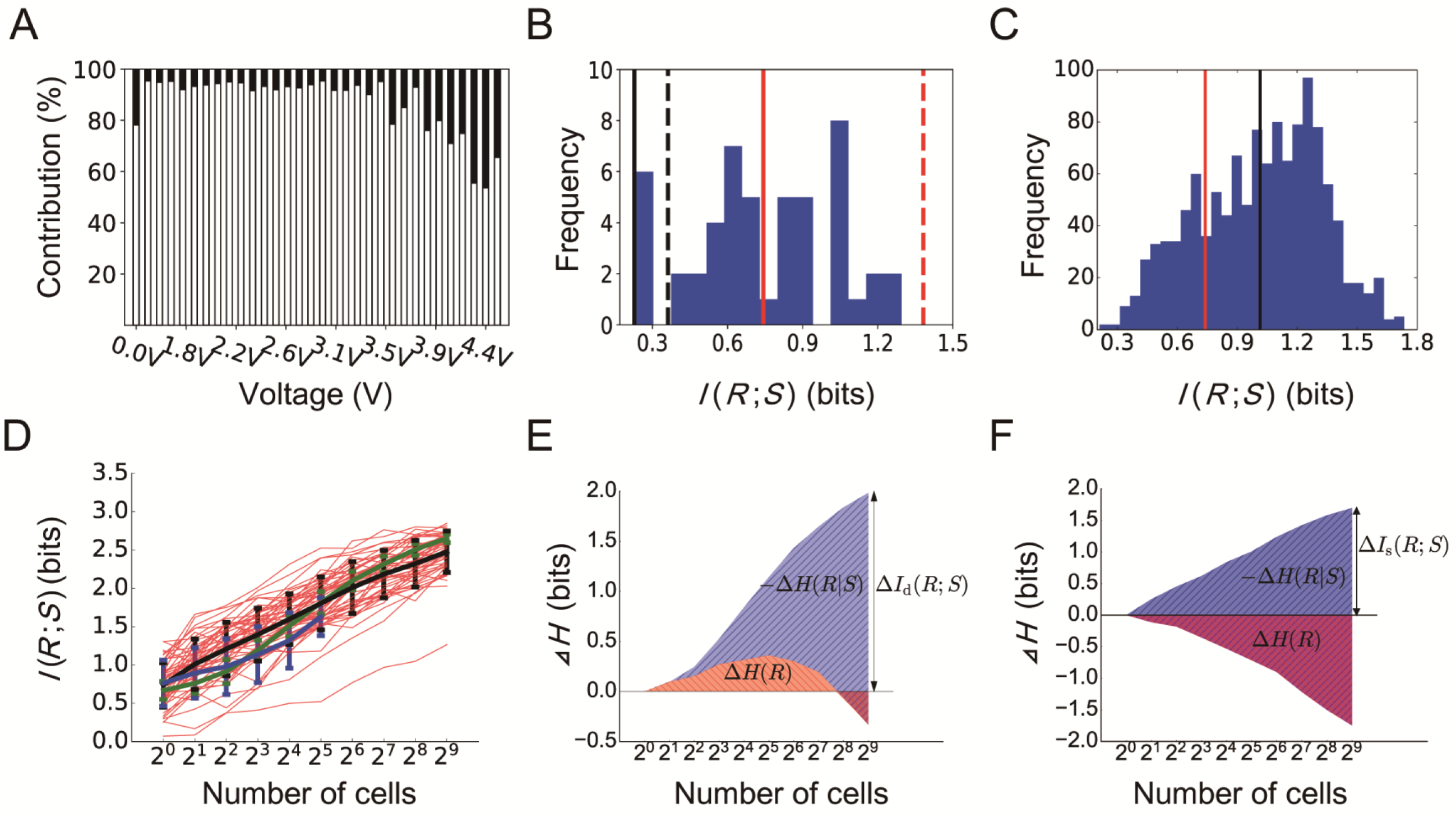
Information transmission of a single-cell channel and a multiple-cell channel for electrical pulse stimulation into contraction of single skeletal muscle fibers. (**A**) The percentages of intercellular (white) and intracellular (black) variation in the total variation for the indicated voltage of EPS in single fibers (Eqs. 2 and 3 in Materials and Methods). The average intracellular variation was 86% of the total variation across all voltages. (**B**) Distribution of the mutual information between intensity of EPS and contraction in single-cell channels. Red dashed line, the average channel capacity of a single-cell channel (1.38 bits); red solid line, the average mutual information of a single-cell channel (0.74 bits); black dashed line, channel capacity of a cell-population channel (0.36 bits); black solid line, mutual information of a cell-population channel (0.25 bits). (**C**) Histogram of the mutual information of multiple-cell channel comprise of 2-cell channels (Eq. 10 in Materials and Methods). Black line, the average mutual information of a 2-cell channel (1.01 bits); red line, the average mutual information of a single-cell channel (0.74 bits). (**D**) Mutual information of a multiple-cell channel according to the number of cells included as single-cell channels. Blue line, the average mutual information of a multiple-cell channel composed of different cells, *I*_d_(*R*; *S*). Bar indicates standard deviation. Red lines, the mutual information of a multiple-cell channel composed of 50 copies of the same cell by resampling responses repetitively from the same cell, *I*_s_(*R*; *S*). Black line, the average *I*_s_(*R*; *S*). Bar indicates standard deviation. Green line, mutual information of an extrapolated multiple-cell channel comprised of different cells generated by extrapolating the standard deviation to the number of cells (Eq. 11 in Materials and Methods). (**E**) The contribution of Δ*H*(*R*) and −Δ*H*(*R*|*S*) to Δ*I*_d_(*R*; *S*) in a multiple-cell channel composed of different single-cell channels. (Eqs. 15, 16 and 17 in Materials and Methods). (**F**) The contribution of Δ*H*(*R*) and −Δ*H*(*R*|*S*) to Δ*I*_s_(*R*; *S*) in a multiple-cell channel composed of copies of the same single-cell channel. We used the optimal input probability distribution for the averaged response of the total cells to calculate the mutual information and the optimal input probability distribution for each channel to calculate channel capacity (dashed lines in B).

As we did for the C2C12 myotubes, we calculated the averaged response of all of the fibers (a total of 50) and calculated the optimal input probability distribution of the averaged response of the total cells, which gives a channel capacity (fig. S5, C and D). We used this optimal input probability distribution to the averaged response of the total cells to calculate the mutual information between EPS and contraction in a single-cell channel for each cell (Fig. 4B), which yielded a value of 0.74 ± 0.29 bits (mean ± S.D.) (Fig. 4B, red solid line). The mutual information of contraction is smaller than that of the Ca^2+^ amplitude in C2C12 myotubes (1.21 bits). Because of the experimental conditions and outputs are different, the mutual information of contraction of single-fiber cells and that of Ca^2+^ amplitude in C2C12 myotubes are not directly comparable, thus we cannot conclude that Ca^2+^ amplitude encodes more information than contraction. Using the optimal input probability distribution for the averaged response of the total fibers, we calculated the mutual information in the cell-population channel and found that the mutual information of the cell-population channel was 0.25 bits (Fig. 4B, black solid line), which is smaller than that in a single-cell channel. Similar to the mutual information between voltage and C2C12 myotube Ca^2+^ response, the mutual information between voltage and fiber contraction is greater when evaluated as a single-cell channel than as a cell-population.

Using the optimal input probability distribution for the averaged response of the total fibers, we calculated the mutual information of 2-cell channels for each pair of fibers. The average mutual information of the 2-cell channel was 1.01 ± 0.32 bits (mean ± S.D.) (Fig. 4C, black, Eq. 10 in Materials and Methods). This result indicates that on average 2.02 conditions can be distinguished by a pair of fibers and that a 2-cell channel can transmit more information than a single-cell channel for which only 1.67 conditions can be distinguished on average. As the number of the fibers composing the multiple-cell channel increased, the mutual information of the multiple-cell channel increased (Fig. 4D, blue), consistent with the increased information capacity that we observed by increasing the number of the cells in the multiple-cell channel for the C2C12 myotubes. Using the same method that we used for the C2C12 myotube data (Fig. 3B), we generated *I*_s_(*R*; *S*) for 50 fibers (Fig. 4D, red). For a 2-cell channel, about 40% of *I*_s_(*R*; *S*) was larger than the average of *I*_d_(*R*; *S*). However, unlike the C2C12 myotube multiple-cell channel, when the number of cells in a multiple-cell channel increased, the percent of *I*_s_(*R*; *S*) (Fig. 4D, red) that exceeded the average *I*_d_(*R*; *S*) (Fig. 4D, blue) did not become smaller. However, the sample size of the single fibers (50) is not as large as that of C2C12 myotubes (551), thus we cannot conclude that *I*_d_(*R*; *S*) does not become larger than *I*_s_(*R*; *S*) with a larger number of cells. Therefore, we extrapolated the standard deviation of an *n*-cell averaged response to the larger number of cells and calculated an *I*_d_(*R*; *S*) using this *n*-cell response (Fig. 4D, green, Eq. 11 in Materials and Methods). This *n*-cell calculation showed the expected reduction in the percent of *I*_s_(*R*; *S*) (Fig. 4D, red) that exceeded the average of the extrapolated *I*_d_(*R*; *S*) (Fig. 4D, green). Thus, the muscle fiber system also transmitted more information as the number of cells of the multiple-cell channel increased as a result of the incorporation of intercellular variation.

We examined the contribution of Δ*H*(*R*) and −Δ*H*(*R*|*S*) to the increase in mutual information of a multiple-cell channel (Fig. 4, E and F, Eqs 13, 14, 16 and 17 in Materials and Methods). The results were the same as for the C2C12 myotube system: Both Δ*H*(*R*) and −Δ*H*(*R*|*S*) contributed to the extrapolated Δ*I*_d_(*R*; *S*) (Fig. 4E), whereas only −Δ*H*(*R*|*S*) contributed to Δ*I*_s_(*R*; *S*) (Fig. 4F). Thus, for both fiber contraction and the Ca^2+^ response in C2C12 myotubes, intercellular variation increases information capacity of a multiple-cell channel.

Intriguingly, there are ~750 cells in flexor digitorum brevis muscle (FDB) of CD-1 mice (*24*). Although we used single fibers from FDB of C57/BL6 mice, we expect that the number of cells in FDB is likely similar in both strains of mice. At 2^9^ (512) cells, the extrapolated *I*_d_(*R*; *S*) was 2.56 bits, which was larger than the average of *I*_s_(*R*; *S*), which was 2.50 bits, suggesting that intercellular variation contributes to accurate contraction of a skeletal muscle.

In this study, we show that a single-cell channel can transmit more information than cell-population channel both for EPS-mediated stimulation of Ca^2+^ signaling in C2C12 myotubes and EPS-mediated contraction of individual skeletal muscle fibers (Fig. 5). Incorporating single-cell channels into a multiple-cell channel has the greatest information transmission capacity. Intracellular variation was smaller than intercellular variation for both biological systems. Intracellular variation is thought to arise from the stochastic fluctuation of biochemical reactions, and intercellular variation is thought to mainly depend on the differences in gene expression and protein abundance among individual cells (*2*). Therefore, smaller intracellular variation means that the fluctuations of biochemical reactions are relatively smaller than the differences produced by gene expression and protein abundance. The larger intercellular variation enables a multiple-cell channel to encode more information than a cell-population or single-cell channel. Taken together, a cellular response is accurate in each cell but different among individual cells, and this difference encodes information in a biological system. Thus, our findings show that “small” intracellular and “large” intercellular variations would enable tissues to precisely respond to a range of stimuli to control physiological function.

**Fig. 5.**
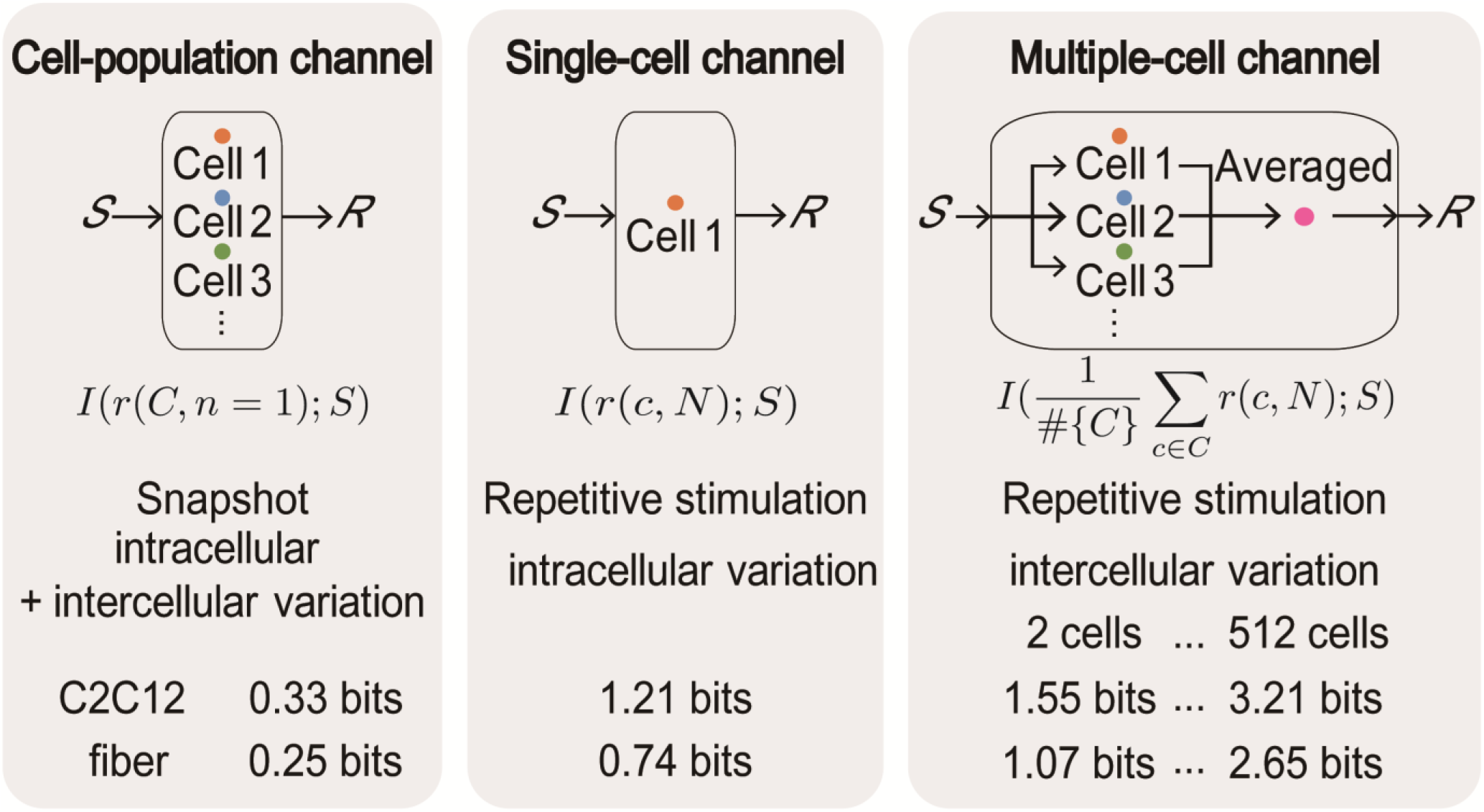
Summary of information capacity in this study. Mutual information of a cell-population channel, the average of mutual information of a single-cell channel and a multiple-cell channel in this study are shown._

In a skeletal muscle, fibers are innervated by a single motor neuron, and a group of fibers and its innervating same motor neuron is called a motor unit (*25*). This means that fibers in the same motor unit receive the same input, whereas fibers in different motor units receive different inputs. Motor neurons vary according to several properties, such as activation threshold and firing rate, resulting in a skeletal muscle contraction by summing contraction of each motor unit, known as recruitment (*26*). As more motor units are recruited, the muscle contraction becomes stronger. In addition to control of contraction by recruitment of motor units, our data indicated that, even fibers with the same input, similar to a motor unit innervated by the same motor neuron, enables precise control of contraction through small intracellular and large intercellular variations. Taken together, our results suggested that control of a motor unit through distinct small intracellular and large intercellular variations in the fibers stimulated by the motor neuron may enable precise control of skeletal muscle contraction.

## Acknowledgments

We thank our laboratory members for critically reading this manuscript and for their technical assistance with the experiments. This manuscript was edited by Nancy R. Gough (BioSerendipity, LLC).

## Author contributions

M.W., T.W., M.K., D.H., K.K., Y.F., Y.M., and N.L.F. carried out the experiments; H.I., T.W., M.W., and M.F. carried out image analysis; T.W., M.W., M.F. S.U. and H.K. carried out computational analysis; writing group consisted of T.W., M.F., M.K., and S.K.; and the study was conceived and supervised by T.W. and S.K.

## Data and materials

Data are available online at http://kurodalab.bs.s.u-tokyo.ac.jp/ja/publication/info/wada/.

## Supplementary Materials

Materials and Methods

Figures S1-S10

Tables S1

