## Supplementary_Materials for "Single-cell information analysis reveals small intra- and large intercellular variations increase cellular information capacity"

**This PDF file includes:**

Materials and Methods
Figs. S1 to S10
Tables S1

### Materials and Methods

Cell culture and the establishment of C2C12 cell lines stably expressing GCaMP

C2C12 cells (kindly provided by Takeaki Ozawa, University of Tokyo, Tokyo, Japan) were cultured in Dulbecco's Modified Eagle's Medium (DMEM; 25 mM glucose; Wako, Japan) supplemented with 10% fetal bovine serum (Nichirei Bioscience Incorporated, Japan) in an incubator at 37°C under a 5% CO2 atmosphere.

C2C12 cells stably expressing the $Ca^{2+}$ biosensor, GCaMP6f (addgene #40755) (*23*), were established with the PiggyBac Transposase System (System Biosciences, U.S.A.) (*27*). One hundred and fifty µL of Opti-MEM (Life technologies, U.S.A.) containing 4 µL of Lipofectamin 2000 (Invitrogen, U.S.A.), and Opti-MEM containing 1.0 µg of PiggyBac transposon vector clone and 0.2 µg of PiggyBac transposase expression vector were mixed and incubated for 5 min. Thereafter, 50% confluent C2C12 cells were seeded on a 35-mm dish, transfected with the mixture, and incubated for 6 h. For the selection of transfected cells, the cells were cultured with DMEM with 10% fetal bovine serum and 20 µg/mL of Blasticidin S Hydrochloride (Wako, Japan). Selected cells were seeded on a Cell Culture Dish (Corning Incorporated, U.S.A.) and cultured until colonies formed. The colonies were picked and seeded on a Cell Culture Dish 430167. After seeding and proliferation, the cells were stored at a concentration of $1.0 \times 10^5$ cells/mL with Bambanker (NIPPON Genetics, Japan).

Differentiation induction of C2C12 cells

C2C12 cells were seeded into 4-well rectangular plates (Thermo Fisher Scientific, U.S.A.) at a density of $2.0 \times 10^5$ cells/well, with 3 mL of 25 mM glucose DMEM supplemented with 10% fetal bovine serum. After 2 days, the medium was switched to a differentiation medium consisting of 25 mM glucose DMEM supplemented with 2% horse serum (Nichirei Bioscience Incorporated, Japan). Ten days after differentiation, cells were used for electrical pulse stimulation (EPS) (*28*).

Fluorescence Microscopy and Electrical pulse stimulation (EPS) to C2C12

Differentiated C2C12 myotubes stably expressing GCaMP6f were washed 3 times with PBS and starved in 3 mL fresh Medium 199, Hanks' Balanced Salts (Life technologies, U.S.A.) for 1 hour. Mineral oil (Sigma Aldrich, U.S.A.) was stratified to prevent vaporization of the medium prior to the fluorescence imaging. Fluorescence imaging was performed with an inverted fluorescence microscope, IX 83 (Olympus, Japan) equipped with a UPLSAPO10X2 objective lens (Olympus, Japan) an ORCA-R2 C10600-10B CCD camera (Hamamatsu Photonics, Japan) , a U-HGLGPS mercury lamp (Molecular Devices, U.S.A.), a U-FBNA mirror unit (Olympus, Japan), an MD-XY30100T-META automatically programmable stage position (Molecular Devices, U.S.A.). EPS was performed according to the method of Manabe *et al.* (*22*). The 4-well plates were connected to the electrical stimulation apparatus, a 4-well C-Dish (Ion Optix Corp., Milton, MA, USA), and stimulated by electric pulses generated by an electrical pulse generator (Uchida Denshi, Hachioji, Japan). Under the microscope, C2C12 myotubes were stimulated with electric pulses of various voltages for 3 ms every 10 seconds and the 70 time-lapse images for each of the 551 cells were acquired every 25 ms for each stimulation. Cells were repetitively stimulated for 20 times each of 10 different voltages, 0, 5, 10, 15, 20, 30, 40, 50, 60, 75 V. A single cycle was composed of one of each of the 10 voltages and each cycle was repeated 20 times, randomizing the order of stimulation independently every cycle. The timing of EPS was controlled with DIO-0808TY-USB digital I/O terminal (CONTEC, Japan) and EPS was imposed 400 ms after opening the shutter of the camera (fig. S1).

Quantification of $Ca^{2+}$ response

After background subtraction from $Ca^{2+}$ time-lapse images acquired by fluorescent microscopy, regions of interest (ROIs) in C2C12 myotubes were selected manually. The time course data for the $Ca^{2+}$ response of each C2C12 myotube were acquired by averaging intensity within each ROI for each image. Basal $Ca^{2+}$ was defined by time average of 16 time points before any stimulation, and $Ca^{2+}$ amplitude was defined by the maximum $Ca^{2+}$ response subtracted by the basal $Ca^{2+}$. AUC was defined by the sum of the difference between $Ca^{2+}$ response and the basal $Ca^{2+}$, multiplied by time step 25 ms.

Single-fiber isolation from a skeletal muscle

Satellite cells were prepared as described previously with modification (*29*). Briefly, 12 week-old C57/BL6J mice were sacrificed by cervical dislocation and the flexor digitorum brevis muscle (FDB) was gently removed. Dissected muscles were then digested in 0.25% collagenase solution (Collagenase Type1; Worthington, Lakewood, NJ) consisting of GlutaMAX Dulbecco's Modified Eagle's Medium (DMEM) supplemented with 1% penicillin-streptomycin and 10% fetal bovine serum (Corning, NY, USA) at 37 °C for 150 min. All tubes, dish, and pipets were coated with 5% bovine serum albumin (BSA) solution in PBS to avoid the adhesion of fibers. FDB was then transferred to a 50-mm Petri dish containing 8 mL of DMEM solution supplemented with 1% penicillin-streptomycin. Under a stereoscopic microscope, the muscles were gently disassembled using 1 mL pipettes to separate them into individual muscle fibers. Cell debris was completely removed by exchanging the solution for fresh DMEM solution numerous times, and 50 fibers were transferred to a 1.5 mL centrifuge tube. The tubes were upright for 5 min to allow the fibers to settle to the bottom, the supernatant was gently aspirated using a Pasteur pipette, and 1 mL of fresh DMEM supplemented with 1% penicillin-streptomycin was added. All of the solution containing the fibers was transferred to 2-well Lab-Tek™ II Chamber Slide (Thermo Fisher Scientific, Waltham, USA) containing 1 mL of fresh DMEM supplemented with 1% penicillin-streptomycin.

Microscopy and EPS-dependent contraction of single muscle fibers

Individual fibers of FDB were stimulated with an electric pulse system that we developed previously for the C2C12 or primary myotubes contraction (*22*, *30*). The 2-well Chamber Slide was connected to the electrical stimulation apparatus and a 2-well electrode (Uchida-denshi, Hachioji, Japan), and fibers were stimulated with electric pulses generated by the power supply (Uchida-denshi). A stimulation cycle consisted of 3 ms duration of EPS and 997 ms intervals at 1 Hz for 10 sec. The initial voltage was 0 V, and the voltage was increased by 0.1 V from 1.5 V to 4.5 V. Contraction of the fibers was recorded under the microscope with the objective lens (X4).

Image analysis and quantification of contraction of single skeletal muscle fibers

After selecting the region with only a single fiber, images were smoothed with a Gaussian filter and the edge region was enhanced with a Sobel filter. The edge-enhanced images were binarized by triangle method. Noises, like spots of binarized images, were removed by the opening filter, holes in fiber region were filled, and noise was removed by the opening filter. The selected regions were regarded as single-fiber regions and the time course data for contraction of each fiber were acquired for each image by quantifying the areas in the selected regions. Basal was defined by time average of last 10 time points of the time course data, and maximal contraction was defined by basal subtracted by minimal area of the time course.

Calculation of probability distribution

Using an adaptive partitioning method (*31*), we calculate probability distributions of $Ca^{2+}$ amplitude and fiber contraction from the experimental data at the single-cell level obtained by repetitive stimulation. We calculated the conditional probability distribution of the response for a given stimulation of a single-cell channel with the single-cell data obtained by repetitive stimulation and that of a cell-population channel with the cell-population data obtained in the first stimulation. $S$, $C$, and $N$ represent the set of stimulation, cell, and the number of stimulations, respectively, and the response $r$ is the function of $C$ and $N$ and can be expressed as $r(C, N)$. The conditional probability distribution of the response for a given stimulation is $p(r(C, N)|S)$. When $c \in C$ is the cell variable and $n \in \mathrm{N}$ is the repeat number variable, the conditional probability distribution of a single-cell channel of cell $c$ is $p(r(c, N)|S)$, and the conditional probability distribution of a cell-population channel calculated by a single stimulation is $p(r(C, n = 1)|\, S)$. By applying the adaptive partitioning method for the experimental data of a single cell, we calculated the probability distribution of $p(r(c, N)|\, S)$. For that of the cell population at a single stimulation, we calculated probability distribution of $p(r(C, n = 1)|\, S)$.

Calculation of the mutual information and channel capacity

The mutual information between probability variables $S$ and $R$ is given by

$$I(R; S) = \sum_{s \in S} p(s) \sum_{r \in R} p(r|s) \log_2 \frac{p(r|s)}{p(r)}, \qquad \text{(Eq. 1)}$$

where $S$ is the stimulation and $R$ is the response, and $p(r|s)$ is $p(r(c, N)|\, S)$ or $p(r(C, n = 1)|\, S)$ as calculated from experimental data. The calculation of the mutual information requires an input probability distribution $p(s)$. An input probability distribution cannot be definitively determined in a biological system; however, the optimal input probability distribution for the maximal mutual information, referred to as the channel capacity, in the channel is often used as an input probability distribution. The channel capactity can be estimated by Blahut-Arimoto algorithm (*32*, *33*). In this study, we defined the "mutual information" the mutual information estimated with the optimal input probability distribution for the average response of the total cells, and we defined "channel capacity" as the mutual information estimated with the optimal input probability distribution for each channel.

Intracellular and intercellular variations

The total variation of each input condition can be divided into intracellular and intercellular variations. Intracellular variation is the sum of the variance of the total cells and can be written as follows:

$$intracellular.variation(s) = \mathrm{E}[\mathrm{Var}[r(c, n)|c, s]|s]. \qquad \text{(Eq. 2)}$$

Intercellular variation is the sum of squares of the difference between the average of each cell and the average of all and can be written as follows:

$$intercellular.variation(s) = \mathrm{Var}[\mathrm{E}[r(c, n)|c, s]|s]. \qquad \text{(Eq. 3)}$$

The total variation was defined by the sum of the intracellular and intercellular variations, and the contributions of intracellular and intercellular variations to the total variation were calculated.

Estimation of the bias in the mutual information calculation caused by sample size, a number of stimulation events at a given dose

We estimated the bias in the calculation of the mutual information using a Hill equation response model (fig. S4). The average dose response was fixed and assumed that to obey the Hill equation,

$$r = \frac{s^n}{s^n + K_D^n}, \quad \text{(Eq. 4)}$$

where *s* is the stimulation variable, *r* is the response variable, *n* is the Hill coefficient and $K_D$ is the dissociation constant. In the toy example shown in fig. S4, $K_D$ is 30 and *n* is 4. We examined the bias caused by small sample size (in our experiments, the number of repetitions of the EPS at each voltage) by adding the Gaussian noise with various standard deviation. On this model, the response is expressed as follows:

$$r = \frac{s^n}{s^n + K_D^n} + \epsilon, p(\epsilon) = \frac{1}{\sqrt{2\pi\sigma^2}} e^{\left(-\frac{\epsilon^2}{2\sigma^2}\right)}, \quad \text{(Eq. 5)}$$

where *s* is the stimulation variable, *r* is the response variable, $\epsilon$ is Gaussian noise with a mean of 0 and the standard deviation is $\sigma$. Gaussian noise is assumed to be independent and have the same variance for all input. On the assumption of continuously and uniformly distributed input from 0 to 75, the averaged dose response was numerically integrated with respect to input so that interdose variation of response was calculated to 0.143. When $\alpha$ is defined as the ratio of intradose variation and the total variation, $\alpha$ is written by the expression:

$$\alpha = \frac{intradose.variation}{intradose.variation + interdose.variation}. \quad \text{(Eq. 6)}$$

So, intradose.variation can be expressed as follows

$$intradose.variation = interdose.variation \frac{\alpha}{1-\alpha}, \quad \text{(Eq. 7)}$$

When $\alpha$ is 0.02, 0.04, 0.08, 0.16, 0.32, 0.64 and 0.9, intradose variation is 0.054, 0.077, 0.112, 0.165, 0.260, 0.504, 1.135, respectively. These intradose variations were used for the standard deviation of Gaussian noise of the Hill model. We sampled various sizes of cells (3, 5, 10, 20, 40, 80, 200, 400, 800, 1000 cells) 30 times for each sample size from the Hill model and estimated the mutual information. We assumed that sample size 1000 is sufficiently large and has ignorable bias. Thus, the calculation bias was defined by the difference between the average mutual information of sample size 20, the number of repetitive stimulations at each voltage that we used in the experiment, and that of sample size 1000.

To examine how many cells there were in the experimental data with $\alpha$ smaller than that used in the model, intradose and interdose variations, and $\alpha$ of each cell were calculated as follows;

$$intradose.variation(c) = \mathrm{E}[\mathrm{Var}[r(c,n)|c,s]|c], \quad \text{(Eq. 8)}$$

$$interdose.variation(c) = \mathrm{Var}[\mathrm{E}[r(c,n)|c,s]|c]. \quad \text{(Eq. 9)}$$

Calculation of the mutual information of a multiple-cell channel

We calculated the mutual information of a multiple-cell channel by calculating the probability distribution of the average responses of multiple cells as follows:

$$p\left(\frac{1}{\#\{\mathrm{C}\}} \sum_{\mathrm{c}\in\mathrm{C}} y(\mathrm{c}, N)\right) | S). \quad \text{(Eq. 10)}$$

The optimal input probability distribution for the average response of the total cells was used to calculate the mutual information of multiple-cell channels. For a 2-cell channel, we calculated the mutual information for all combinations of pairs of the single cells. When the responses of 2-cells were combined, we used all combinations of the responses to the same stimulation (Fig 3A and 4C).

To investigate the contribution of the intracellular and intercellular variations to the increase in the mutual information achieved by combining single-cell channels, we calculated the mutual information of a multiple-cell channel composed of the same cell $I_{\mathrm{s}}(R;S)$ (Fig. 3B and 4D, red lines) and the mutual information of a multiple-cell channel composed of different cells $I_{\mathrm{d}}(R;S)$ (Fig. 3B and 4D, blue line). To calculate $I_{\mathrm{s}}(R;S)$, we virtually created the multiple-cell channel by resampling responses 1, 2, 4, 8, 16, 32, 64, 128, 256, 512 times repetitively to the same stimulation from the same cell, randomizing the order of responses every time. $I_{\mathrm{s}}(R;S)$ was defined as the mutual information of the average response of the virtual multiple-cells and was calculated once for each cell for each resample size. We calculated $I_{\mathrm{d}}(R;S)$ by resampling 1, 2, 4, 8, 16, 32, 64, 128, 256, 512 cells for the $Ca^{2+}$ amplitude in C2C12 myotubes, and 1, 2, 4, 8, 16, 32 cells for the contraction of single fibers with replacement from the total cells. $I_{\mathrm{d}}(R;S)$ was defined as the mutual information of the average response of the resampled multiple cells and calculated 100 times for each resample size. The resampling was performed independently for each resample size. The order of responses was randomized every time the response was resampled.

Because there were only 50 cells for single fibers, we extrapolated the standard deviation to 1, 2, 4, 8, 16, 32, 64, 128, 256, 512 cells and calculated an extrapolated $I_{\mathrm{d}}(R;S)$ (Fig. 4D, green line). We assumed that the average of dose response reaches equilibrium and does not change over 50 cells, and the standard deviation becomes smaller as the number of cells increases. The noise obeys Gaussian distribution and is independent for all input. When *n* is the number of cells, the standard deviation of *n*-cell averaged response is

$$\sigma(n) = \sigma(50) * \sqrt{\frac{50}{n}}. \qquad \text{(Eq. 11)}$$

<u>The contribution of $\Delta H(R)$ and $\Delta H(R|S)$ to $\Delta I(R;S)$</u>

The mutual information can be expressed by the entropy of response $H(R)$, and the conditional entropy of response for a given stimulation $H(R|S)$ as follows:

$$I(R;S) = H(R) - H(R|S). \qquad \text{(Eq. 12)}$$

With the input probability distribution and the calculated response probability distribution, $H(R)$ and $H(R|S)$ can be calculated as follows:

$$H(R) = \sum_{s\in S} p(s) \sum_{r\in R} p(r|s) \log_2 p(r), \qquad \text{(Eq. 13)}$$

$$H(R|S) = \sum_{s\in S} p(s) \sum_{r\in R} p(r|s) \log_2 p(r|s). \qquad \text{(Eq. 14)}$$

The difference of the mutual information of two different channels, defined as $\Delta I(R;S)$, can be written as

$$\Delta I(R;S) = \Delta H(R) - \Delta H(R|S), \qquad \text{(Eq. 15)}$$

where $\Delta H(R)$ is the difference of the entropy of response, and $\Delta H(R|S)$ is the difference of the conditional entropy of response for a given stimulation. The differences were defined by

$$\Delta H(R) = H_n(R) - H_1(R), \qquad \text{(Eq. 16)}$$

$$\Delta H(R|S) = H_n(R|S) - H_1(R|S), \qquad \text{(Eq. 17)}$$

where $H_n(R)$ and $H_n(R|S)$ represent the average of $H(R)$ and $H(R|S)$ of 100 resampled populations of *n*-cell channels (Fig. 3, C and D, and 4, E and F, fig. S9).

<u>Binary channel with Gaussian noise</u>

We built the binary response model, with a fixed threshold in each cell but different among individual cells (fig. S9). The response of each cell can be written as

$$r(s) = \begin{cases} 1 + \epsilon \ (\text{if } s \geq threshold) \\ 0 \ + \epsilon \ (\text{if } s < threshold) \end{cases}, \qquad \text{(Eq. 18)}$$

where $s$ is the stimulation variable, $r$ is the response variable, $\epsilon$ is Gaussian noise with a standard deviation of 0.1. The threshold is fixed in each cell but different among individual cells and is continuously and uniformly distributed from 0 to 1 in the cell population. We assumed two kinds of input probability distribution: One is continuous uniform distribution from 0 to 1, and the other is discrete uniform distribution of 0, 0.1, 0.2, 0.3, 0.4, 0.5, 0.6, 0.7, 0.8, 0.9, 1.0. We resampled 1, 2, 4, 8, 16, 32, 64, 128, 256, 512 cells 100 times from this model and calculated the average of $I_{\mathrm{d}}(R;S)$ for each number of cells when continuous and discrete input probability distribution was given.

**Table S1. A summary of previous studies that calculated mutual information of signaling pathways.** The mutual information in a cell-population channel calculated by a single stimulation, in a single-cell channel, and a multiple-cell channel in previous and this study are summarized.

| Input | Output | MI (bits) | channel | sample size per dose | Literature |
|---|---|---|---|---|---|
| Bcd | Hb | 1.5 ±0.15 (mean ± S.D.) | Cell-population | < 1000 | Tkačik *et al.* (*5*) |
| TNF | NF-κB, ATF-2 | 0.92, 0.85 | Cell-population | 350 | Cheong *et al.* (*12*) |
| NGF,PACAP, PMA | pERK,pCREB, c-Fos,EGR1 | > 1 | Cell-population | 1000～2000 | Uda *et al.* (*14*) |
| EGF, ATP, LPS | ERK, $Ca^{2+}$, NF-κB | > 1.7 | Cell-population | < 5320 | Selimkhanov *et al.* (*16*) |
| GnRH | ppERK, EGR1, NFAT-NF, NFAT-RE | > 1 | Cell-population | < 10000 | Garner *et al.* (*17*) |
| ATP | $Ca^{2+}$ | > 1.2 | Cell-population | < 2500 | Potter *et al.* (*18*) |
| Ach | $Ca^{2+}$ | 2.06 ± 0.31 (mean ± S.D.) | Single-cell* | 5 | Keshelava *et al.* (*21*) |
| EPS | <br>$Ca^{2+}$<br>Contraction<br><br><br><br>$Ca^{2+}$<br>Contraction | Single-cell<br>1.21 ± 0.50<br>0.74± 0.29<br>(mean ± S.D.)<br>**Multiple-cell**<br>(512 cells)<br>3.21<br>2.65 | Single-cell<br>**Multiple-cell** | 20 | This study |

*By interpolating the response probability distribution, the mutual information was calculated with continuous input probability distribution. In this study, we did not interpolate the response (fig. S9), the mutual information of a single-cell channel in this study may be underestimated compared to the study by Keshelava *et al.* (*21*).

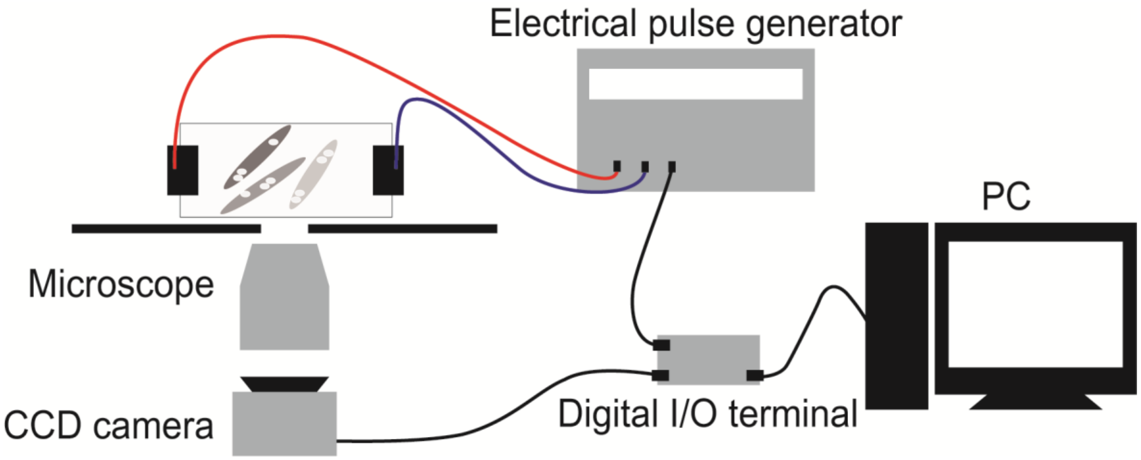


**Fig. S1. Experimental setup of EPS-dependent $Ca^{2+}$ response in C2C12 myotubes and of cell-contraction in single-fiber cells.** When the signal reached the digital I/O terminal from the computer (PC), the terminal sent the signals to the CCD camera and the electrical pulse generator to start both image acquisition and EPS. For C2C12 myotubes, the terminal sent a signal to the camera 400 ms before the signal to the electrical pulse generator. For single-fiber cells, the terminal sent a signal to the camera 50 ms before the signal to the electrical pulse generator. Signal was then sent to the electrical pulse generator every one second.

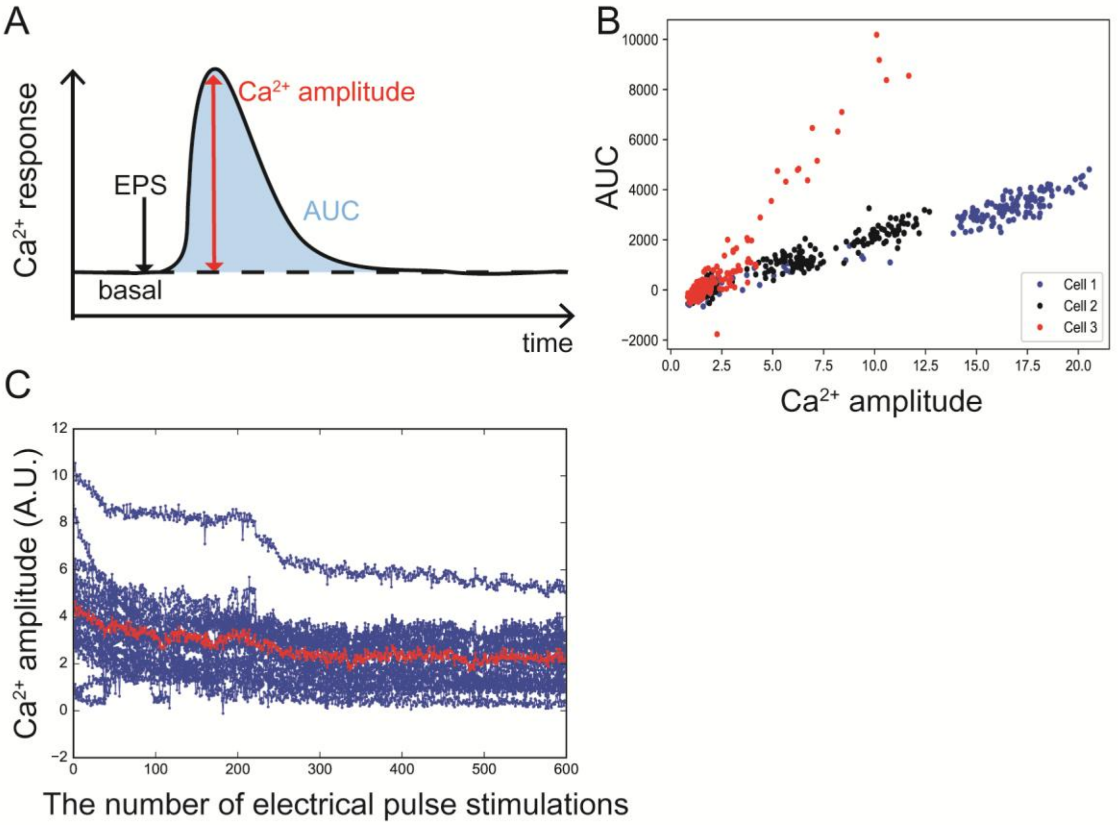


**Fig. S2. The definition of $Ca^{2+}$ amplitude and $Ca^{2+}$ amplitude by repetitive EPS(30V).** (**A**) $Ca^{2+}$ amplitude (red arrow) and AUC (blue area) were defined (Materials and Methods). (**B**) Correlation coefficient between $Ca^{2+}$ amplitude and AUC in each cell was 0.88 ± 0.13 (mean ± S.D.), indicating $Ca^{2+}$ amplitude and AUC were positively correlated. We used $Ca^{2+}$ amplitude as the feature value of $Ca^{2+}$ response. (**C**) $Ca^{2+}$ amplitude by repetitive EPS stimulation (30V). Blue: $Ca^{2+}$ amplitude in each cell ($n$=19). Red: the ensemble average. The amplitudes of many cells remained the same through 200 stimulations. Amplitude decay started when the number of repetitions exceeded 200. We set the total number of repetitions to 200.

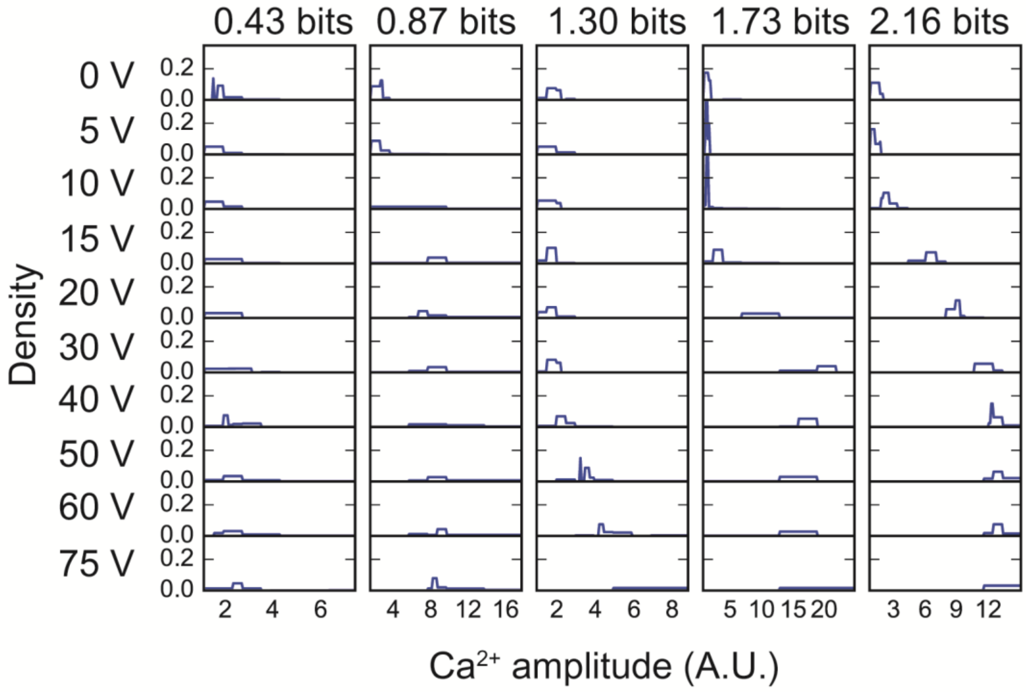


**Fig. S3. The conditional probability distribution of $Ca^{2+}$ amplitudes for a given stimulation of the cells having various mutual information.** In setting the total number of stimulations, the number of inputs and the sample size in each stimulation has a trade-off relationship. Using the 10 conditions, 0, 5, 10, 15, 20, 30, 40, 50, 60, 75 V, we examined the conditional probability distribution of $Ca^{2+}$ amplitude in C2C12 myotubes for a given stimulation. The mutual information for each myotube is indicated at the top and the data are presented in order of increasing mutual information. Because of few leaps of the position of probability distribution between adjacent input conditions for the cells having various different mutual information, we concluded that these 10 conditions were sufficient to avoid underestimation of the mutual information in a single-cell channel.

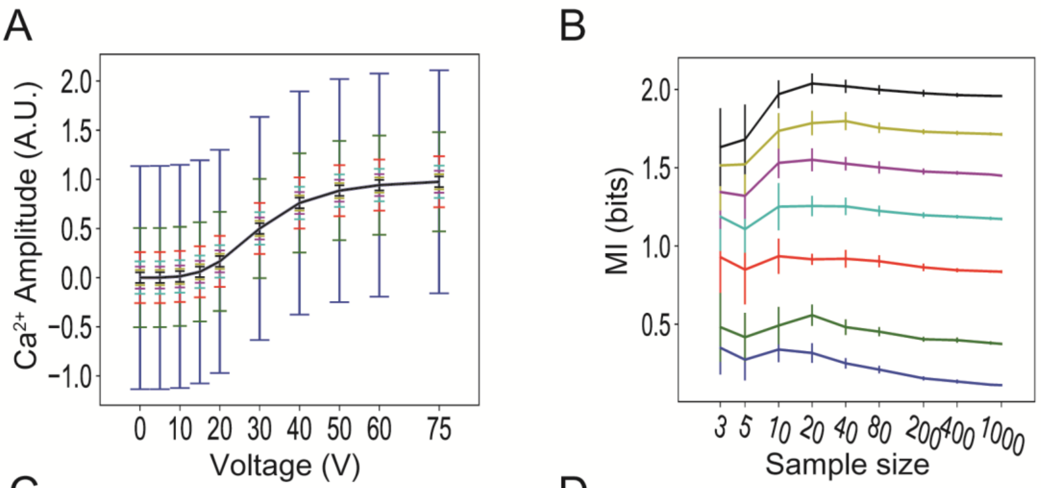


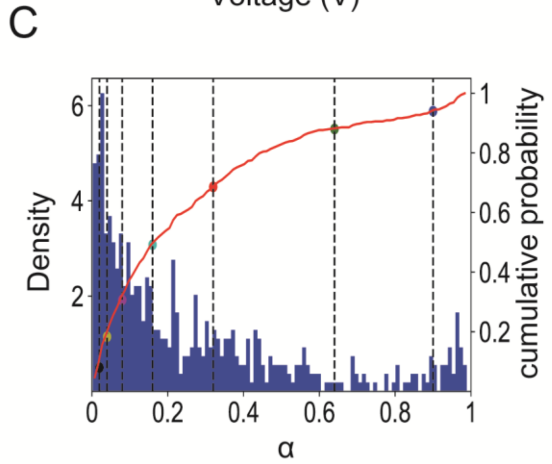


D

| $\alpha$ | $\sigma$ | Percentile | $I_{20}$(bits) mean±SD | $I_{1000}$(bits) mean±SD | $\Delta I$(bits) | $2^{I_{20}} - 2^{I_{1000}}$ |
|---|---|---|---|---|---|---|
| 0.02 | 0.054 | 8 % | 2.04±0.06 | 1.96±0.01 | 0.08 | 0.24 |
| 0.04 | 0.077 | 19 % | 1.78±0.07 | 1.71±0.01 | 0.06 | 0.16 |
| 0.08 | 0.112 | 32 % | 1.54±0.07 | 1.45±0.01 | 0.08 | 0.16 |
| 0.16 | 0.165 | 50 % | 1.28±0.05 | 1.17±0.01 | 0.11 | 0.17 |
| 0.32 | 0.260 | 69 % | 0.91±0.09 | 0.83±0.01 | 0.11 | 0.14 |
| 0.64 | 0.504 | 88 % | 0.54±0.07 | 0.37±0.01 | 0.17 | 0.16 |
| 0.90 | 1.135 | 94 % | 0.33±0.06 | 0.112±0.004 | 0.21 | 0.17 |

**Fig. S4. Estimation of bias caused by sample size, the number of repetitions at each stimulation, calculated with a toy mathematical model.** (**A**) A dose response of the Hill model from a toy mathematical model of Ca2+ amplitude in response to 4 stimulations at each voltage. The model includes Gaussian noise with various standard deviations. In the Hill model, the response to the stimulation can be expressed as $r = \frac{s^n}{s^n + K_D^n} + \epsilon, p(\epsilon) = \frac{1}{\sqrt{2\pi\sigma^2}} e^{\left(-\frac{\epsilon^2}{2\sigma^2}\right)}$ (Eq. 5). In the example, $K_D$ is 30, Hill coefficient $n$ is 4, $s$ is a stimulation variable, $r$ is a response variable, $\epsilon$ is a Gaussian noise with a mean of 0, and the standard deviation is $\sigma$. $\alpha$ is defined as the ratio of intradose variation to the total variation, the sum of intradose and interdose variations (Eq. 6). When the values of $\alpha$ are 2, 4, 8, 16, 32, 64, and 90%, the values of intradose variation $\sigma$ are 0.054, 0.077, 0.112, 0.165, 0.260, 0.504, and 1.135, respectively. Gaussian noise is added independently with each $\sigma$ for all doses. Color indicates noise at each $\alpha$: 2% (black), 4% (yellow), 8% (magenta), 16% (cyan), 32% (red), 64% (green), and 90% (blue). (**B**) The relationship of sample size (number of repetitions of the stimulation) and calculated mutual information by adding Gaussian noises with various standard deviations. Bars indicate the standard deviation. (**C**) Histogram of calculated $\alpha$ of each cell from the experimental data (Eqs. 6, 8, and 9). Red line indicates the empirical distribution. Dashed line indicates the given $\alpha$. (**D**) The bias of mutual information when the sample size is 20. The value of $\alpha$ for the models is given, $\sigma$ is the standard deviation calculated from the given $\alpha$, "percentile" is the percentage of the cells with smaller $\alpha$ than the given value. As $\alpha$ becomes smaller, , the smaller $\sigma$ becomes.

$I_{20}$ and $I_{1000}$ are the mutual information when sample sizes are 20 and 1000, respectively. $\Delta I = I_{20} - I_{1000}$ is the bias in the mutual information, and $2^{I_{20}} - 2^{I_{1000}}$ is the bias in the average number of controllable states. At a sample size of 20, representing the number of repetitive stimulations used in the experiments, bias in the number of controllable states was an overestimation ranging from 0.14 to 0.24. Because our data with the C2C12 myotubes resulted in 2.32 controllable states for a single-cell channel and 1.70 controllable states for a cell-population channel (determined from the optimal input), even the maximal bias (when $\alpha$ was 2%) does not affect the conclusion that a single-cell channel can transmit more information than a cell-population channel. Moreover, only ~8% of the myotubes had an $\alpha$ less than 2%. Therefore, we concluded that calculation bias existed but had little effect on results and interpretation and that the sample size of 20 repetitive stimulations at each voltage was adequate.

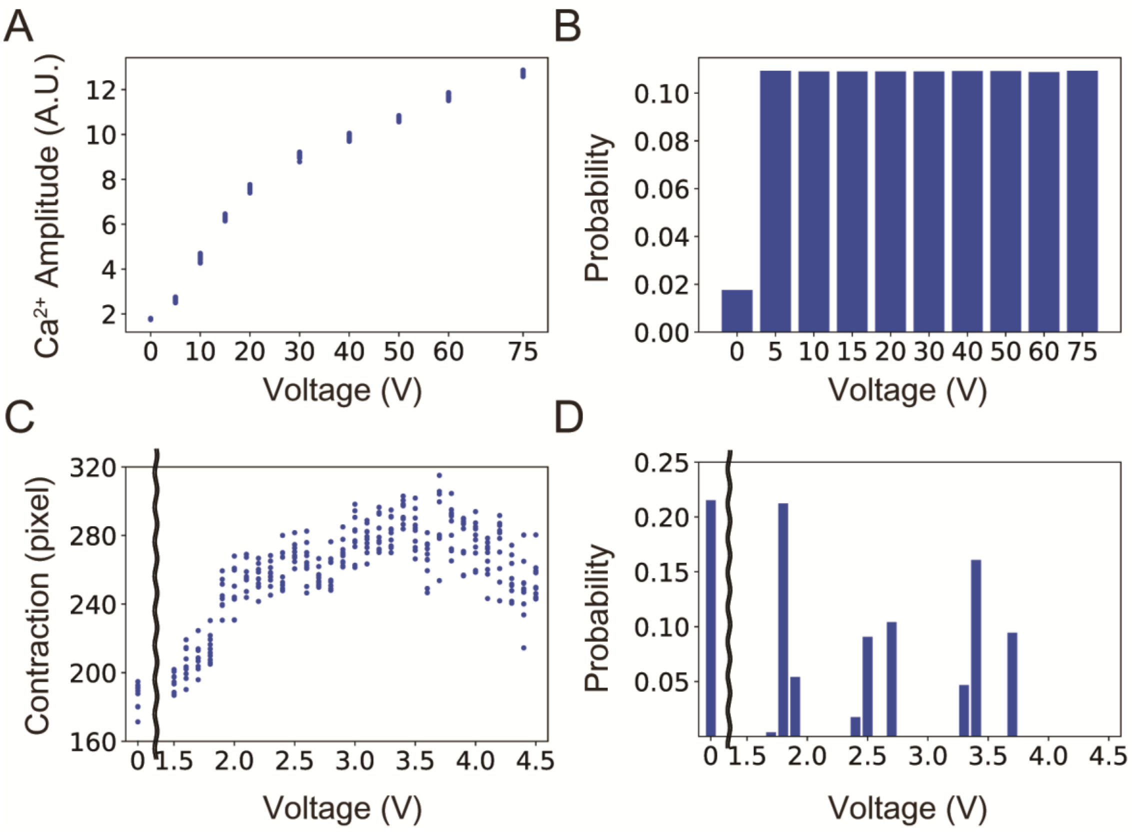


**Fig. S5. The average response of the total cells and the optimal input probability distribution for the average response.** (**A**) The averaged response of the $Ca^{2+}$ amplitudes at each voltage (0 – 75 V) for all 551 C2C12 myotubes. (**B**) The optimal input probability distribution for the average response of the $Ca^{2+}$ amplitude in all 551 C2C12 myotubes. (**C**) The average response of contraction for all 50 skeletal muscle fibers stimulated from 0 – 4.5 V.(**D**) The optimal input probability distribution for the average response of contraction of all 50 skeletal muscle fibers.

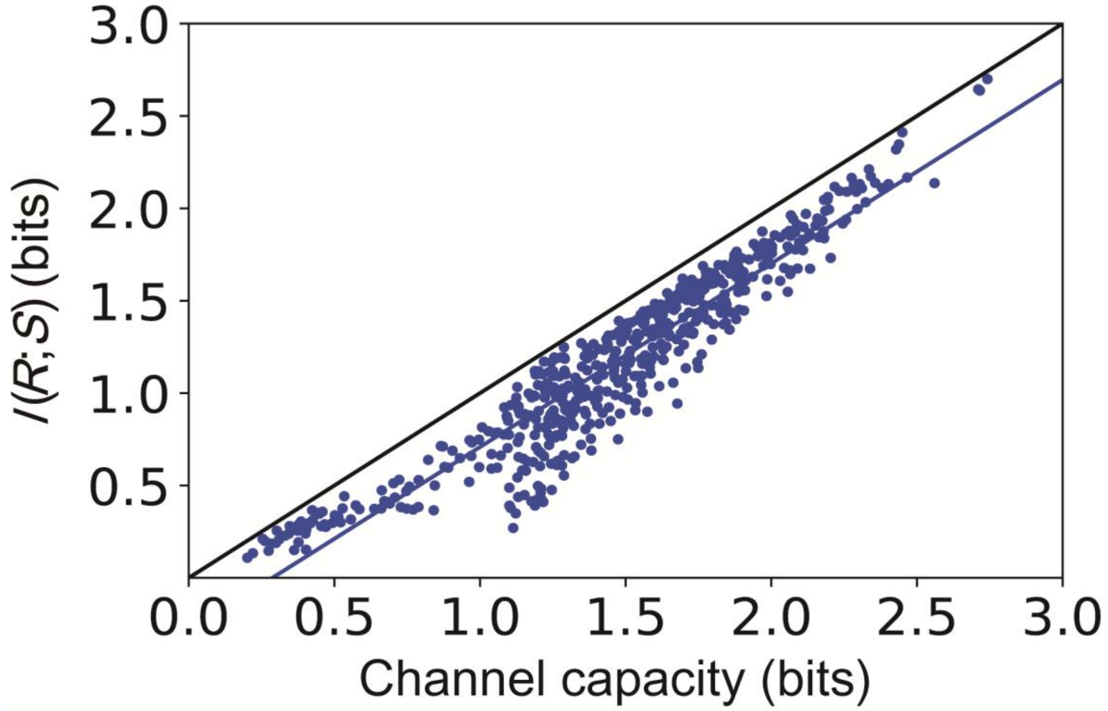


**Fig. S6. Mutual information with the optimal input probability distribution for the average response and channel capacity of single-cell channels in C2C12 myotubes.** A dot indicates the mutual information calculated with the optimal input probability distribution for the average response and the channel capacity of single-cell channels. The black line indicates $y = x$; the blue line indicates the regression line ($y = 0.993\ x - 0.285$ with correlation coefficient, 0.947). We used the optimal input probability distribution for the average response of the total cells to calculate the mutual information, and we used the optimal input probability distribution for each single-cell channel to calculate the channel capacity.

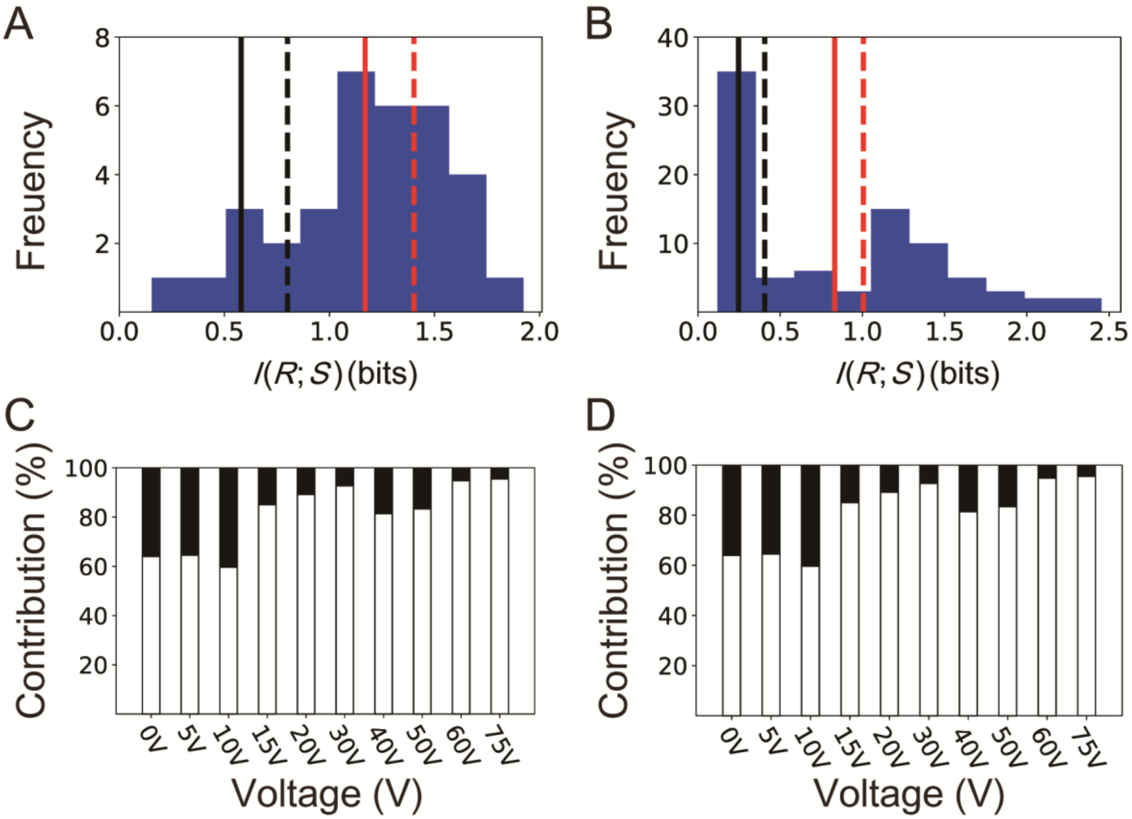


**Fig. S7. Mutual information in two other independent clones (#2, #3) of C2C12 myotubes stably expressing GCaMP6f.** (**A** and **B**) Histograms of mutual information of single-cell channels in clone #2 (A) and #3 (B). The numbers of cells of clone #2 and #3 were 34 and 86 respectively. Red solid line, the average of the mutual information of single-cell channels (1.19 bits for clone #2; 0.85 bits for clone #3); red dashed line, the channel capacities of single-cell channels (1.43 bits for clone #2; 1.02 bits for clone #3); black solid line, channel capacities of cell–population channels (0.75 bits for clone #2; 0.38 bits for clone #3); black dashed line, the mutual information of cell–population channels (0.55 bits for clone #2; 0.26 bits for clone #3 (B). In both clones, the mutual information of the average of single-cell channels was larger than that of a cell-population channel. (**C** and **D**) The percentages of intercellular (white) and intracellular (black) variations in the total variation by the indicated voltage of EPS in a clone #2 (C) and #3 (D) (Eqs. 2 and 3). In both clones, intercellular variations were larger than intracellular variations. We used the optimal input probability distribution for the average response of the total cells to calculate the mutual information, and we used the optimal input probability distribution for each channel to calculate the channel capacity (fig. S7 A, B, dashed lines).

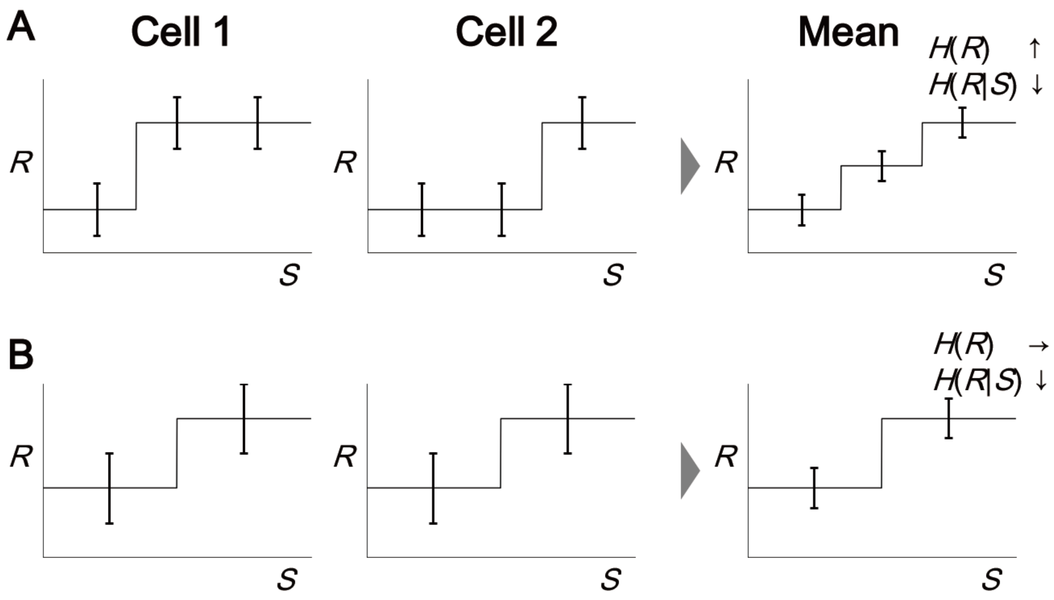


**Fig. S8. The effect of dose responses of 2 cells with different or identical responses on mutual information of a 2-cell channel.** (**A**) When a pair of cells have different dose responses, $H(R)$ can increase because of an increase in the distinguishable number by averaging two different dose responses. Averaging the multiple-cell responses decreases $H(R|S)$, because the variance of the average response decreases as the number of cells increases. Thus, both the increase in $H(R)$ and the decrease in $H(R|S)$ contribute to the increase in the mutual information. (**B**) When a pair of cells has the identical dose response, $H(R)$ does not increase, because averaging the dose responses does not result in a dose response that is different from that of either of the two cells. Averaging the multiple-cell responses decreases $H(R|S)$. Thus, only the decrease in $H(R|S)$ contributes to the increase in the mutual information.

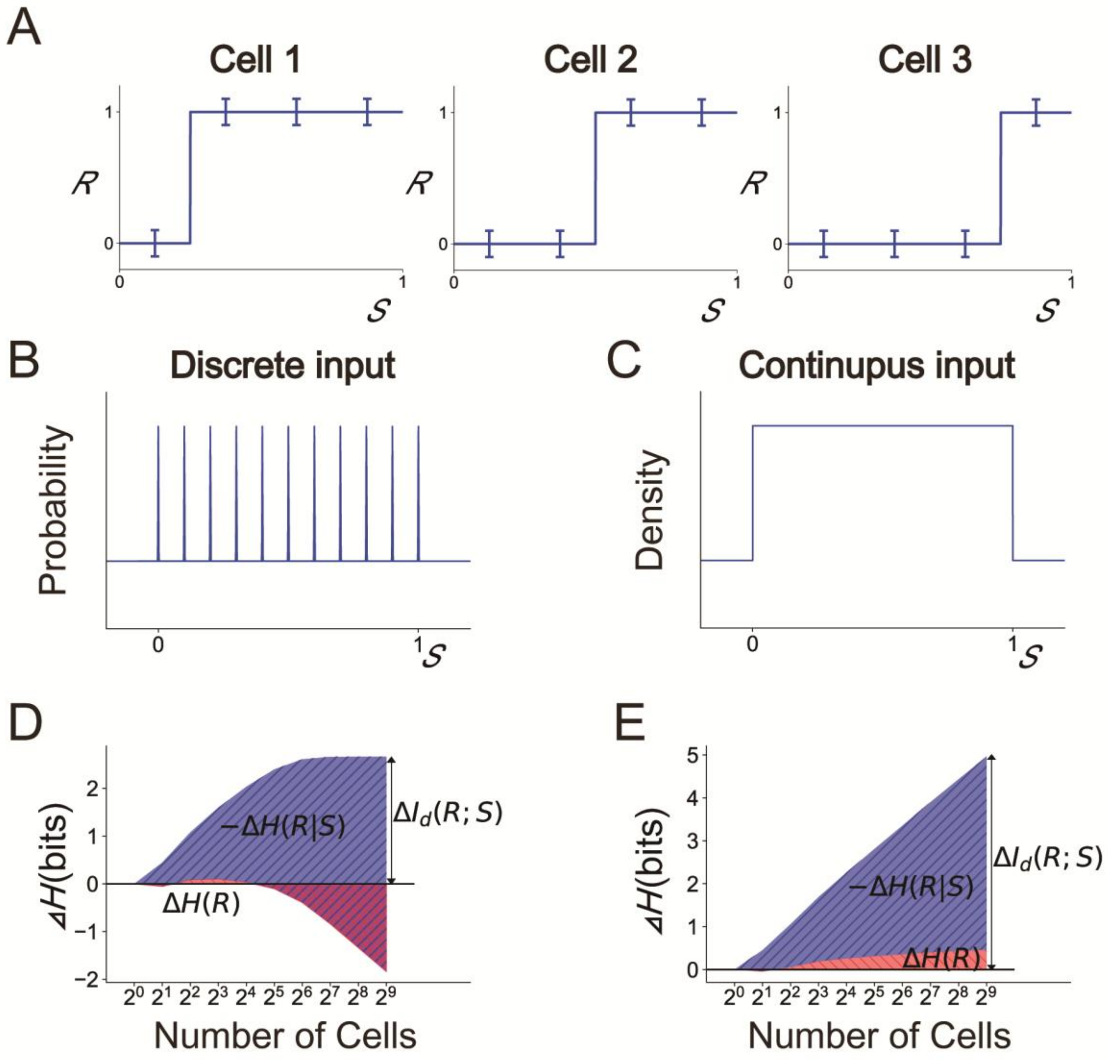


**Fig. S9. The effect of discrete and continuous input on $\Delta H(R)$ and $-\Delta H(R|S)$ in a binary response model.** (**A**) A binary response model, with a threshold that is different among individual cells and evenly distributed from 0 to 1 (Eq. 18) (Materials and Methods). Each cell responds 0 if $s$ is smaller than the threshold and responds 1 if $s$ is larger than the threshold. Gaussian noise with a standard deviation of 0.1 was added independently to the total cells and to the stimulation. (**B**) Discrete uniform input probability distribution of 0, 0.1, 0.2, 0.3, 0.4, 0.5, 0.6, 0.7, 0.8, 0.9, 1.0. (**C**) Continuous uniform input probability distribution from 0 to 1. (**D**) The contribution of $\Delta H(R)$ and $-\Delta H(R|S)$ to the mutual information in a multiple-cell channel composed of different single-cell channels, $\Delta I_{\mathrm{d}}(R;S)$, with discrete input probability distribution. As the number of cells increases, $\Delta H(R)$ increases and then decreases. The differences $\Delta I_{\mathrm{d}}(R;S)$,

$\Delta H(R)$, and $\Delta H(R|S)$ for each number of cells were defined by the differences from those when the number of cells is 1 (Eqs. 15, 16, 17). (**E**) The contribution of $\Delta H(R)$ and $-\Delta H(R|S)$ to the mutual information in a multiple-cell channel composed of different single-cell channels, $\Delta I_{\mathrm{d}}(R;S)$, with continuous input probability distribution. As the number of cells increases, $\Delta H(R)$ increases monotonically. The differences $\Delta I_{\mathrm{d}}(R;S)$, $\Delta H(R)$, and $\Delta H(R|S)$ for each number of cells were defined by the differences from those when the number of cells is 1 (Eqs. 15, 16, 17).

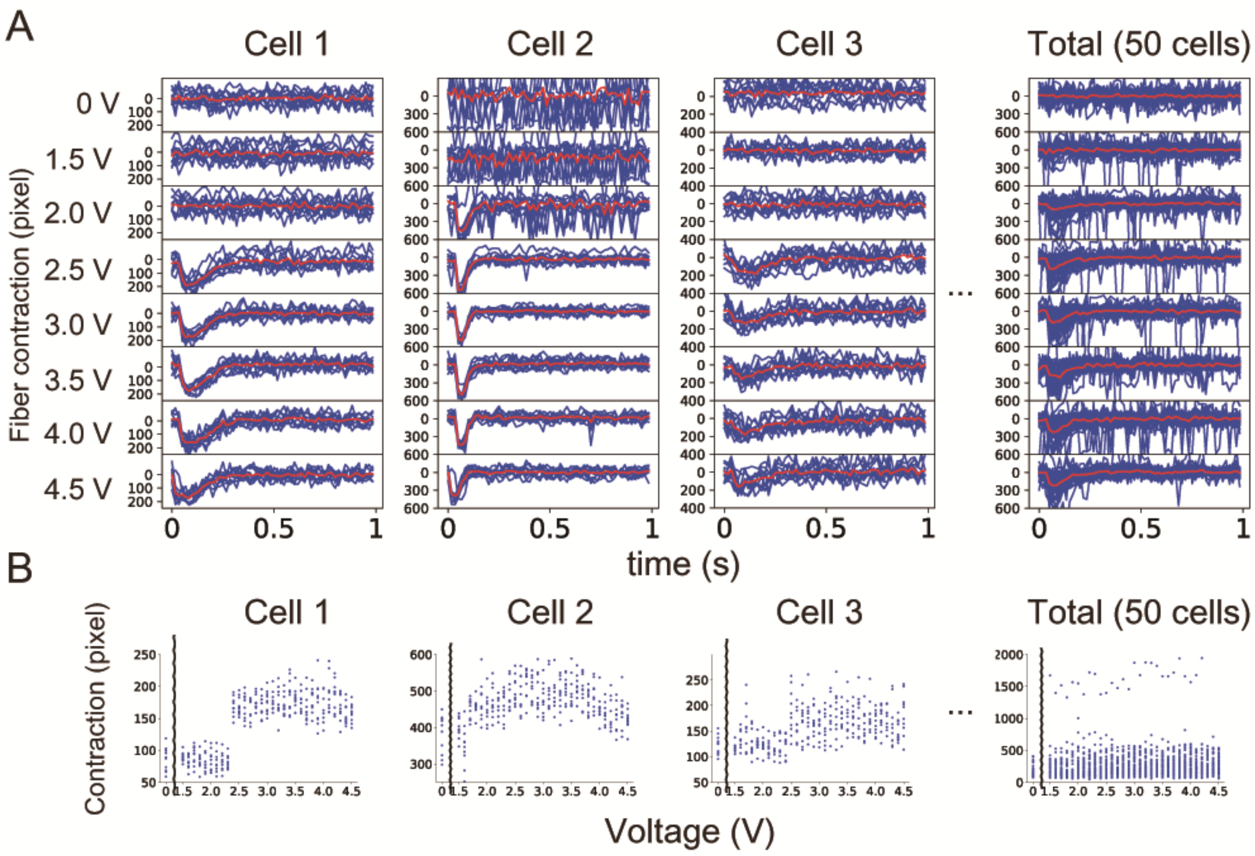


**Fig. S10. Contraction of single muscle fibers.** (**A**) Contraction in individual fibers in response to 10 times-repetitive stimulation with 32 different voltages of EPS from 0 – 4.5 V. "Cell 1" to "Cell 3" are representative of the contraction of single fibers. "Total" indicates responses in 50 fibers. Each blue line indicates a time course of the contraction by a single stimulation for each cell. Red lines indicate the averaged time course. (**B**) Dose responses of contraction for the data shown in (A). A dot indicates maximal contraction induced by eac h single stimulation.
